# Brain signaling dynamics after vagus nerve stimulation

**DOI:** 10.1101/2021.06.28.450171

**Authors:** Vanessa Teckentrup, Marina Krylova, Hamidreza Jamalabadi, Sandra Neubert, Monja P. Neuser, Renée Hartig, Andreas J. Fallgatter, Martin Walter, Nils B. Kroemer

**Affiliations:** University of Tübingen, Department of Psychiatry and Psychotherapy, Tübingen, Germany; Max Planck Institute for Biological Cybernetics, Tübingen, Germany; IMPRS for Cognitive and Systems Neuroscience, Tübingen, Germany; University of Magdeburg, Department of Psychiatry and Psychotherapy, Germany; Leibniz Institute for Neurobiology, Magdeburg, Germany; Department of Psychiatry, University Hospital Jena, Jena, Germany

## Abstract

The vagus nerve projects to a well-defined neural circuit via the nucleus tractus solitarii (NTS) and its stimulation elicits a wide range of metabolic, neuromodulatory, and behavioral effects. Transcutaneous vagus nerve stimulation (tVNS) has been established as a promising technique to non-invasively alter brain function. However, the precise dynamics elicited by tVNS in humans are still largely unknown. Here, we performed fMRI with concurrent right-sided tVNS (vs. sham) following a randomized cross-over design (*N*=40). First, to unravel the temporal profile of tVNS-induced changes in the NTS, we compared fMRI time series to canonical profiles for stimulation ON and OFF cycles. Model comparisons indicated that NTS time series were best fit by block-wise shifts in signal amplitude with stimulation ON and OFF estimates being highly correlated. Therefore, we compared stimulation (ON + OFF) versus baseline phases and found that tVNS increased fMRI BOLD activation in the NTS, but this effect was dependent on sufficient temporal signal-to-noise ratio (tSNR) in the mask. Second, to identify the spatiotemporal evolution of tVNS-induced changes in the brain, we examined lagged co-activation patterns and phase coherence. In contrast to our hypothesis, tVNS did not alter dynamic functional connectivity after correction for multiple comparisons. Third, to establish a positive control for future research, we measured changes in gastric myoelectrical frequency via an electrogastrogram. Again, in contrast to our hypothesis, tVNS induced no changes in gastric frequency. Collectively, our study provides evidence that tVNS can perturb brain signaling in the NTS, but these effects are dependent on tSNR and require precise localization. In light of an absence of acute tVNS-induced effects on dynamic functional connectivity and gastric motility, we discuss which steps are necessary to advance future research on afferent and efferent effects of tVNS.

## 1. Introduction

To survive, any organism must adjust its motivated behavior, such as food seeking, to recuperate its energetic demands. To this end, the brain receives feedback from peripheral organs, for example, the gut and the stomach, on the current metabolic state of the body (Gribble & Reimann, 2019; Small & DiFeliceantonio, 2019). This information is largely transmitted via the vagus nerve, a key part of the autonomic nervous system projecting to the nucleus tractus solitarii (NTS; de Lartigue, 2016; Howland, 2014). These afferent vagal signals are sufficient to limit food intake as decerebrated rats can still terminate their intake in response to caloric load (de Lartigue, 2016; Grill & Norgren, 1978). Thus, a close correspondence between physiological signals reflecting metabolic need and the brain is vital to achieve energy homeostasis. Consequently, metabolic disorders, such as obesity, have been linked to disturbances within this circuit (Cork, 2018). Whereas there is emerging evidence on the role of vagal afferent signals in the control of motivated behavior (Davis et al., 2008; de Lartigue, Ronveaux, & Raybould, 2014; Tellez et al., 2013), it is still largely elusive how vagal input shapes whole-brain signaling dynamics underlying behavioral control.

Despite the confined anatomical target of vagal afferents, their stimulation has been associated with a variety of neural, neuromodulatory, and behavioral effects. For example, it is well established that the NTS relays metabolic information to the mid- and forebrain (de Lartigue, 2016; Grill & Hayes, 2012). In humans, implanted cervical vagus nerve stimulation has been shown to increase cerebral blood flow in the dorsal anterior cingulate cortex and the dorsal striatum (Conway et al., 2012, 2006). Furthermore, vagal afferents have been shown to modulate dopaminergic (Han et al., 2018; Tellez et al., 2013), noradrenergic (Roosevelt, Smith, Clough, Jensen, & Browning, 2006), and cholinergic signaling (Hulsey et al., 2016). Endogenous stimulation of the vagus nerve is induced by nutrients and also evokes dopamine responses in the dorsal striatum tracking caloric value (de Araujo, Ferreira, Tellez, Ren, & Yeckel, 2012; Ferreira, Tellez, Ren, Yeckel, & de Araujo, 2012). Behaviorally, vagal afferent signals modulate appetite (Bodenlos et al., 2007) and food intake (de Lartigue et al., 2014; Val-Laillet, Biraben, Randuineau, & Malbert, 2010). Nevertheless, behavioral effects extend beyond food, as episodic and spatial memory (Suarez et al., 2018), cognitive flexibility (Klarer, Weber-Stadlbauer, Arnold, Langhans, & Meyer, 2017), mood (Klarer et al., 2014), and reinforcement learning (Kühnel et al., 2020) were found to be affected by vagal modulation as well. Taken together, this body of evidence suggests that vagal afferent signals play an important neuromodulatory role, although it is not clear yet how the effects in the brain can be most effectively induced in humans.

Recently, non-invasive transcutaneous vagus nerve stimulation (tVNS) has become a promising novel technique to modulate vagal afferents along a well-defined neuroanatomical pathway. tVNS capitalizes on the dense innervation of the inner ear by the auricular branch of the vagus nerve (Howland, 2014). Only in the past decades, experimental studies have established the feasibility of non-invasive electrical vagal stimulation by demonstrating far-field potentials from the brain stem after tVNS (Fallgatter et al., 2003). Critically, tVNS does not substantially alter heart rate or blood pressure that might confound hemodynamics (Kraus et al., 2007).Thus, tVNS can be used concurrently to fMRI. Neuroimaging studies have demonstrated increased activation in areas well in line with afferent vagal projections throughout the brain stem (i.e., NTS, substantia nigra, dorsal raphe, locus coeruleus, and periaqueductal gray; Dietrich et al., 2008; Frangos, Ellrich, & Komisaruk, 2015; Yakunina, Kim, & Nam, 2017). Changes in brain activation after tVNS have also been found in neuroanatomically connected brain regions (Table 1) suggesting that tVNS elicits brain activation changes within an extended NTS circuit. This regulatory circuit is vital for energy homeostasis, as Yao et al. (2018) recently demonstrated that an implanted closed-loop VNS system effectively reduces food intake and delays weight gain in rats. Likewise, non-invasive tVNS has been shown to affect gastric motility by causing increased amplitude and decreased frequencies of action potentials in human gastric muscle cells (Hong et al., 2018). These tVNS-induced changes in stomach movements and brain activation might be directly linked via the “gastric network”, a resting-state network synchronized with the gastric rhythm (Rebollo, Devauchelle, Béranger, & Tallon-Baudry, 2018). Since the gut-brain circuit can be targeted via tVNS, characterizing alterations within this network could provide crucial insights into signaling dynamics potentially leading to optimized stimulation protocols in humans.

**Table 1.** Prior functional magnetic resonance imaging studies reporting brain areas that are neuroanatomically connected to brain stem nuclei and showed activation changes after transcutaneous vagus nerve stimulation

| Change in brain activation | Study |
| --- | --- |
| + NTS | (Frangos et al., 2015) (S) [mostly ipsilateral] |
|  | (Frangos & Komisaruk, 2017) (S) [ipsilateral] |
|  | (Yakunina, Kim, & Nam, 2017) (S) [bilateral] |
|  | (Sclocco et al., 2019) (BL) [bilateral] |
| - NTS | (Frangos & Komisaruk, 2017) (S) [contralateral] |
|  | (Kraus et al., 2013) (S) [mostly bilateral] |
| + Parabrachial nuclei | (Frangos et al., 2015) (S) |
|  | (Frangos & Komisaruk, 2017) (BL, S) |
| + Thalamus | (Dietrich et al., 2008) (BL) |
|  | (Kraus et al., 2007) (S) |
|  | (Kraus et al., 2013) (S) |
|  | (Yakunina, Kim, & Nam, 2017) (S) |
| + Hypothalamus | (Yakunina, Kim, & Nam, 2017) (S) |
| - Hypothalamus | (Frangos et al., 2015) (S) |
| - Visual cortex | (Frangos & Komisaruk, 2017) (BL) |
| + Amygdala | (Frangos et al., 2015) (S) |
| - Amygdala | (Kraus et al., 2007) (S) |
| + Caudate | (Yakunina, Kim, & Nam, 2017) (S) |
|  | (Badran et al., 2018) (S) |
| + Insula | (Dietrich et al., 2008) (BL) |
|  | (Frangos et al., 2015) (BL) |
|  | (Frangos & Komisaruk, 2017) (BL, S) |
|  | (Kraus et al., 2013) (S) |
|  | (Kraus et al., 2007) (S) |
| + Basal ganglia | (Frangos & Komisaruk, 2017) (BL, S) |
|  | (Yakunina, Kim, & Nam, 2017) (S) |
| - Temporal gyrus | (Kraus et al., 2013) (S); (Kraus et al., 2007) (S) |
| - Hippocampus | (Kraus et al., 2007) (S) |
|  | (Frangos et al., 2015) (S) |
|  | (Frangos & Komisaruk, 2017) (BL, S) |
| - Parahippocampal gyrus | (Kraus et al., 2007) (S)<br>(Kraus et al., 2013) (S) |
| + Nucleus accumbens<br>- Nucleus accumbens | (Frangos et al., 2015) (S)<br>(Dietrich et al., 2008) (BL) |
| + Paracentral lobule | (Frangos et al., 2015) (S) |
| + Postcentral gyrus | (Dietrich et al., 2008) (BL) |
| + Precentral gyrus | (Kraus et al., 2007) (S) |
| + Primary sensory cortex | (Frangos & Komisaruk, 2017) (BL) |
| + Cingulate gyrus | (Badran et al., 2018) (S)<br>(Dietrich et al., 2008) (BL) |
| + Frontal cortex | (Badran et al., 2018) (S)<br>(Dietrich et al., 2008) (BL)<br>(Frangos & Komisaruk, 2017) (BL, S) |
| - Posterior cingulate cortex | (Kraus et al., 2013) (S) |

(+) Increase, (-) Decrease, (BL) compared to baseline, (S) compared to sham stimulation

To develop targeted clinical applications in humans, detailed insights into circuit-level brain dynamics are decisive (Berényi, Belluscio, Mao, & Buzsáki, 2012; Dayan, Censor, Buch, Sandrini, & Cohen, 2013; Kringelbach, Green, & Aziz, 2011; Muldoon et al., 2016; Saenger et al., 2017), and tVNS provides an experimental means to causally alter signaling cascades in a homeostatic network. Arguably, causal manipulation of brain signaling can be achieved with other stimulation techniques, such as transcranial magnetic stimulation (TMS), transcranial direct current stimulation (tDCS) or deep brain stimulation (DBS) as well. While TMS uses electromagnetic induction to de- or hyperpolarize cortical neurons (Walsh & Pascual-Leone, 2003), tDCS can be used to temporarily shift the resting membrane potential of cortical neurons (Utz, Dimova, Oppenländer, & Kerkhoff, 2010). Both techniques, though, can only be used to modulate cortical areas close to the skull and lack the anatomical specificity of tVNS (Beam, Borckardt, Reeves, & George, 2009; George & Aston-Jones, 2010; Horvath, Forte, & Carter, 2015; Klooster et al., 2016). Moreover, the efficacy of tDCS with conventional settings is still controversially debated in the literature (Boayue et al., 2019; Horvath et al., 2015; Opitz et al., 2016) and there is currently no positive outcome control available for tDCS. In contrast, for TMS, induced motor events in the abductor pollicis brevis can be used as positive control for depolarization within the motor cortex (Mogg et al. 2007, Pridmore et al. 1998). Higher anatomical specificity is provided by DBS, which allows stimulation of neural tissue deep in the brain (Kringelbach, Jenkinson, Owen, & Aziz, 2007). To verify the position of the electrodes, electrophysiological measures can be used during surgery (Milosevic et al., 2018; Neumann et al., 2019; Vitek, Delong, Starr, Hariz, & Metman, 2011). In combination with lead externalization after surgery, these measures can be used as positive control (Neumann et al., 2019; Rosa et al., 2017; Voges, Müller, Bogerts, Münte, & Heinze, 2013). However, due to the invasive nature of DBS, research is restricted to therapeutic applications precluding use in healthy participants (Cusin, Soskin, & Dougherty, 2012). Taken together, in contrast to other brain stimulation techniques, tVNS may provide a non-invasive way to perturb a well-defined brain circuit to investigate causally induced signaling cascades in the brain, but a positive control is still missing.

### Hypotheses

tVNS might serve as a non-invasive and spatially well-defined technique to perturb neural circuits related to metabolism. Thus, the overarching goal of this study is to characterize changes in brain dynamics induced by tVNS that are currently not well understood in humans. Consequently, we used model comparisons to determine potential temporal profiles of brain activation induced by tVNS. Furthermore, we aimed to unravel the spatiotemporal progression of tVNS effects by employing complementary connectivity indices that track changes elicited by the stimulation in space and time.

#### 1. Temporal evolution of the tVNS effect (main objective)

We expected tVNS to increase activation in the brain stem, particularly in the NTS. We hypothesized that tVNS either leads to 1) a ramping increase, 2) an initial impulse function followed by a linear decrease, or 3) a block-wise change in amplitude. This temporal prediction was based on the default stimulation protocol of the NEMOS tVNS device, consisting of 30s ON and OFF cycles. We tested for the temporal profile of the tVNS effect by generating synthetic fMRI time series by convolving the three-by-three candidate activation “models” with the canonical hemodynamic response function. Then, we fit hierarchical linear models accounting for random effects of participants to predict the fMRI time-series extracted from the NTS region of interest (ROI). To identify the best fitting model of the temporal profile at the first level, we used model comparisons. Moreover, since the impact of participant variables modulating the effects at the second level is still largely elusive, we finally compared two versions of the winning temporal model: One with only order of stimulation conditions as a control variable at the second level, and one with BMI and sex as additional control variables.

#### 2. Spatiotemporal profile of the tVNS effect (secondary objective)

Since tVNS effects are not spatially confined, we hypothesized that the tVNS-induced change in activation causes a cascade propagating downstream (Table 1). To unravel the pathway of this signaling cascade, we tested for regions that show a time-lagged activity which is associated with the primary tVNS effect as assessed within the NTS by Hypothesis 1. To this end, we used complementary connectivity measures that provide point estimates (i.e., for each TR) to avoid averaging over multiple sequential scans. On the one hand, we tested for lagged tVNS effects based on signal *amplitude* by performing incremental calculations of co-activation patterns (CAPs) with increasing lag. On the other hand, we calculated dynamic phase-coherence to determine changes in *frequency* that are orthogonal to changes in amplitude. In other words, dynamic phase-coherence capitalizes on phase similarity within normalized (i.e., free of differences in amplitude) time courses of brain signal.

#### 3. tVNS-induced effects on myoelectrical frequency (positive control outcome)

Based on recent findings, we expected that tVNS reduces gastric frequency (positive control). Thus, to validate successful acute stimulation of the vagal pathway, we measured an electrogastrogram (EGG) and modeled changes in gastric frequency for tVNS versus sham using linear mixed-effects models.

To summarize, although non-invasive tVNS has gained considerable traction in recent years, the dynamics of afferent effects on brain function are still largely unknown. Likewise, an independent positive control to verify tVNS has not been established so far. Thus, we applied tVNS versus sham stimulation in 40 healthy participants using a single-blind randomized cross-over design. First, to uncover the temporal profile of the stimulation effect in the target region NTS, we concurrently measured brain function using fMRI at rest. Second, to uncover the spatio-temporal progression of the stimulation effect, we analyzed time-shifted connectivity maps originating from the NTS. Third, we concurrently acquired the gastric myoelectric frequency via an electrogastrogram to evaluate successful stimulation of the NTS based on efferent modulation of the gut. After improving the SNR of the mask to achieve good coverage of the NTS signal, we found a significant increase in activity with low-to-moderate levels of smoothing, but not in models using unsmoothed time series. In contrast to our hypotheses, we found no effect of tVNS on dynamic functional connectivity (FC) or gastric frequency.

## 2. Methods

### 2.1 Participants

In total, we recruited 45 healthy participants via public announcement, including posts on the email lists of the University of Tübingen and University Hospital Tübingen, social media and flyers. Five participants had to be excluded from the final sample. Three of these participants left the study by request before both sessions had been completed. One participant was excluded due to motion parameters exceeding the preregistered threshold (> 50% of the total number of volumes exceeding a framewise displacement of 0.5 mm) in the resting-state fMRI measurement. One participant was excluded because the stimulation failed to start precisely at the beginning of the stimulation phase of the resting-state fMRI measurement. This yielded the final sample of 40 healthy participants (22 female, M_age_ = 25.5 years ± 6.6; M_BMI_ = 24.0 kg/m² ± 3.1). All participants completed a phone screening prior to taking part in the study. To be eligible, the following criteria had to be met: 1) 18-50 years of age; 2) BMI range of 18.5-30 kg/m²; 3) no lifetime history of brain injury, cardiovascular diseases, schizophrenia, bipolar disorder, epilepsy, diabetes or asthma; 4) no implants (e.g. pacemaker, cochlear implant, cerebral shunt; except dental prostheses); 5) within the last 12 months: no severe substance abuse disorder, anxiety disorder (except specific phobia), obsessive compulsive disorder, trauma- and stressor-related disorder, somatic symptom disorder or eating disorder; 6) no open wounds or impaired skin at electrode site; 7) not pregnant or nursing; 8) be eligible for MR research (i.e., no non-removable metal parts, such as piercings, no tattoos above the neck or larger than 14 cm, no claustrophobia, noise tolerability). All participants provided written informed consent prior to the first session. The protocol has been approved by the ethics committee of the medical faculty of the University of Tübingen and the University Hospital Tübingen (reference number 235/2017BO1).

### 2.2 Power analysis

To provide accurate within-subject estimates of tVNS-induced effects, we aimed to investigate 40 participants. To determine an adequate sample size for the main objective, we ran simulations using the intended design^1^. We based our simulation on block-wise stimulation-induced brain activation, a candidate model which mirrors the NEMOS protocol. Accordingly, we first generated design regressors and convolved them with the canonical hemodynamic response function provided in SPM12 (spm_volterra, TR = 1 s to simplify calculations; see 2.5.3 Statistical analysis, temporal evolution of the tVNS effect). We then simulated BOLD time series for tVNS and sham conditions, respectively, using known parameter values. To generate the tVNS BOLD time series, we randomly drew N “beta” values from a normal distribution (*M* = 0.2, *SD* = 0.12), where N was the number of participants. Next, we multiplied this beta with the design regressors for each participant. Lastly, we added random measurement noise by sampling from a normal distribution (*M* = 0, *SD* = 1). As sham stimulation should lead to less pronounced activation in the ROI, we reasoned that the average beta would be smaller. To account for the nested repeated measures design (i.e., correlations of stimulation effects among participants), beta values for the sham conditions were generated by multiplying each tVNS beta with a beta parameter controlling the reliability of the measurement (*beta*_reliability_ = 1.15) and subsequently subtracting 0.11 from the result. We then added noise drawn randomly from a normal distribution (*M* = 0, *SD* = 0.1). Lastly, we recovered “observed” beta values by estimating them from simulated BOLD time series for tVNS and sham based on the design regressors (regress function as implemented in MATLAB R2017a (The Mathworks Inc., Natick, MA, USA)). To calculate the effect size Cohen’s *d_z_*, we estimated the mean differences between betas (tVNS-sham) and the *SD* of the differences and calculated the ratio.

To get the sampling distribution, we repeated this procedure 1000 times. Accordingly, a sample size of N = 40 allowed us to recover large within-subject effects (median latent Cohen’s *d_z_* ∼1.00 vs. recovered *d_z_* ∼0.79 given a correlation of *r* = .57 between sessions corresponding to moderate test-retest reliability (Fröhner, Teckentrup, Smolka, & Kroemer, 2019)). Thus, for this basic within-subject comparison, the power exceeded 99% and provided excellent precision of the effect-size estimate, 95% CI [0.45, 1.17], which is important for the model comparisons to be conclusive.

### 2.3 Experimental procedure

Our study (randomized single-blind cross-over design) consisted of 2 sessions following the same standardized protocol (Figure 1). Thus, participants received tVNS in one of the sessions, and sham stimulation in the other session with order being determined in advance. We used the function randperm as implemented in Matlab 2018a to generate a shuffled vector of zeros and ones coding for the stimulation condition in the first experimental session with the second session being the complementary condition of the cross-over design (the exported Excel table, locked prior to study commencement, can be accessed on OSF: https://osf.io/uqtke/).

**Figure 1.**
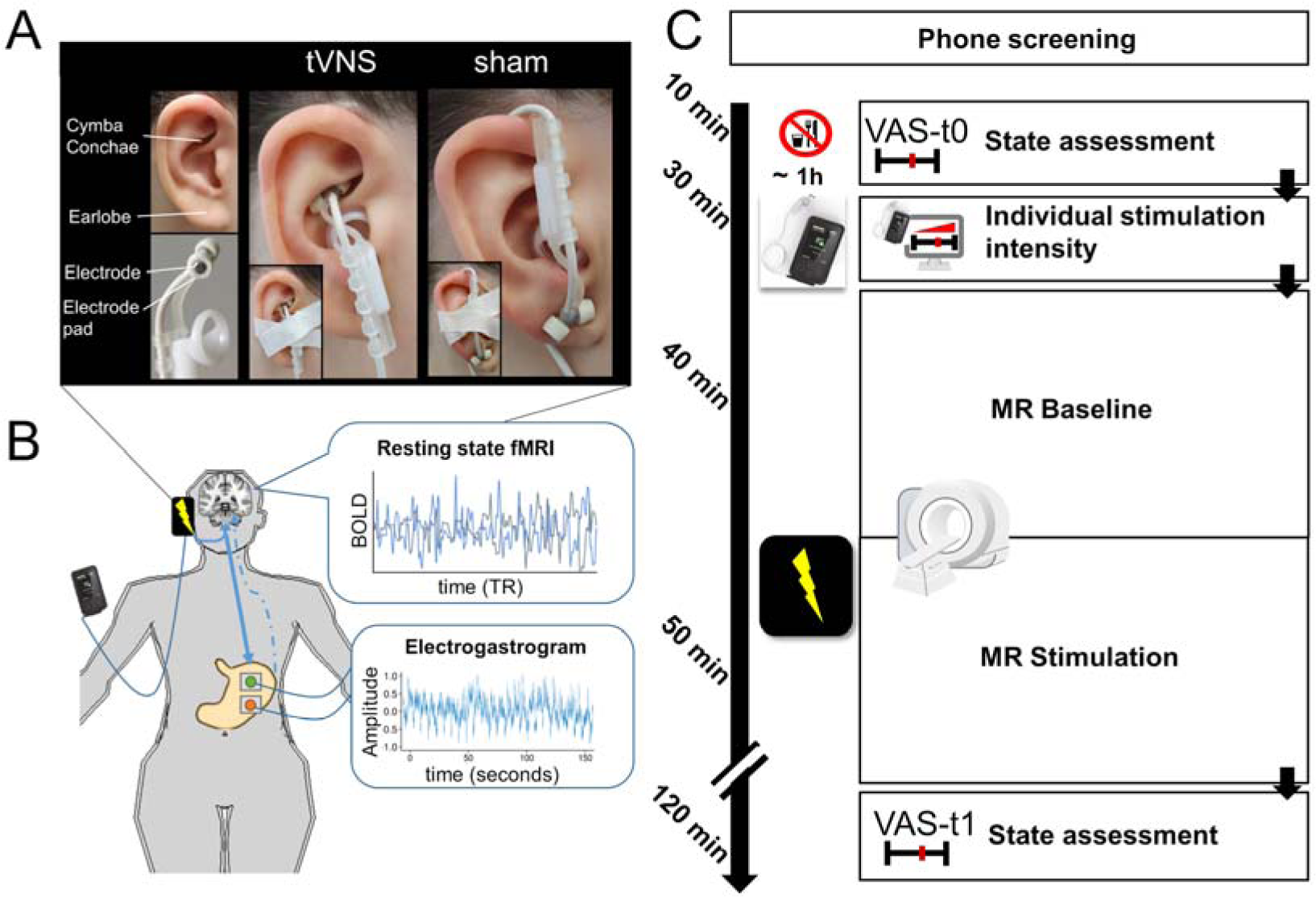
Schematic summary of the experiment. A: Placement of the electrode for transcutaneous vagus nerve stimulation (tVNS) and sham condition. B: Depiction of the targeted pathway and the methods used to investigate afferent and efferent effects of tVNS. C: Procedure for the study. Visualization was generated using R (https://www.r-project.org/) and gganatogram (Maag, 2018).

Participants were asked to enter the experimental session neither hungry nor full and to refrain from consuming food or caloric beverages 1 h prior to each session. They were further asked to eat approximately 1.5 h prior to the beginning of each session to ensure comparable delays to the last meal. At the beginning of each session, anthropometric measures were taken, including body weight, height, and waist and hip circumference (World Health Organization, 2011). Participants also provided information about their last intake of food and drinks and, if applicable, their menstrual cycle and oral contraceptive use. Thereafter, we acquired ratings of hunger, fullness, thirst, and mood based on items of the PANAS (Watson, Clark, & Tellegen, 1988), according to our previously established procedure for tVNS testing (Kühnel et al., 2020).

After participants completed state ratings, the stimulation site was cleaned with ethanol and the stimulating electrodes were placed on the cymba conchae for tVNS and on the earlobe for sham stimulation as described in Frangos et al. (2015). Medical white tape was used to durably attach the electrodes to the desired location. Then, we tested if the electrodes had sufficient skin contact by briefly activating the stimulator. Before participants entered the scanner, EGG and ECG electrodes were positioned. Furthermore, to reduce artifacts due to head motion, we cleaned the participant’s forehead with ethanol and applied a strip of medical tape from one side of the magnetic resonance head coil to the other via the forehead which serves as a tactile feedback if the participant moves (Krause et al., 2019). Once participants were positioned inside the scanner, the EGG baseline measurement started. Then, we determined the stimulation intensity with the electrodes being placed at the sham or tVNS site, respectively, at the beginning of each session based on the individual pain threshold (corresponding to mild pricking). After defining this threshold, the electrode remained in place, but was not active until participants entered the stimulation phase.

During the MR measurement, we first ran a localizer (1 minute) and acquired the T1-weighted anatomical scan (6 minutes). During this anatomical scan, participants completed training rounds for the tasks. Task fMRI was measured within the study, but this data is not a measure of interest for the current report. After acquiring fieldmaps (2 minutes), we showed instructions for the following resting-state fMRI (rs-fMRI) measurement on a screen inside the scanner room, asking the participant to lie still, to think of nothing in particular, and to stay awake. Then, the rs-fMRI baseline was measured (10 minutes). While rs-fMRI scans were running, we showed a 10-minute version of the Inscapes animation (without audio) to improve compliance while minimizing cognitive load and motion (Vanderwal, Kelly, Eilbott, Mayes, & Castellanos, 2015). After 10 minutes of baseline rs-fMRI, stimulation was turned on. Then, 10 minutes of rs-fMRI were collected during active stimulation (tVNS or sham). To ensure precise estimates of myoelectric activity, EGG measurement started before rs-fMRI (after positioning) and continued thereafter while the stimulation was still on.

Thereafter, we continued with two reward processing tasks with concurrent active stimulation (60 minutes). These tasks are no measure of interest for the current report. Afterwards, the stimulation was turned off, the EGG recording was stopped and the participant was taken out of the scanner. After a short break, participants completed the state ratings for a second time (∼5 minutes). At the end of each session, participants received the rewards they earned based on their task performance (extra monetary payment and snacks) and a baseline compensation for participation in the study (either 56 € or partial course credit) at the end of the second session.

### 2.4 tVNS device

Stimulation was applied transcutaneously to the auricular branch of the vagus nerve using Cerbomed NEMOS (Erlangen, Germany), a non-invasive, stimulation device (Kreuzer et al., 2012; Van Leusden, Sellaro, & Colzato, 2015). NEMOS has been established as a therapeutic tool (Bauer et al., 2016) and is compatible with concurrent fMRI measurements (Frangos et al., 2015). Given the current use of this device in therapeutic contexts, investigating the effects of the stimulation protocol on brain dynamics is of high relevance.

With NEMOS, electrical stimulation was applied to the right ear via a titanium electrode placed at the cymba conchae (tVNS) or the same electrode turned upside down and placed at the earlobe (sham), following the protocol of (Frangos et al., 2015; Figure 1). The stimulation protocol of NEMOS is preset with a biphasic impulse frequency of 25 Hz with alternating intervals of 30 s stimulation ON and 30 s OFF. We adjusted the individual stimulation intensity based on subjective pain thresholds using VAS ratings at the beginning of each session (for details, see Kühnel et al. (2020)). In short, we initialized the stimulation at 1 mA and increased stepwise in 0.5 mA units. Participants rated the current sensation (“How intensely do you feel pain induced by the stimulation?”) ranging from 0 (“no sensation”) to 10 (“strongest sensation imaginable”) until they settle around 5 (“mild pricking”).

To safely apply the stimulation inside the MR scanner, the stimulator was placed in the control room. Only the earpiece with the electrodes was located in the scanner room and connected to the stimulator via a cable routed through an RF filter plate.

### 2.5 Magnetic resonance imaging

#### 2.5.1 Data acquisition

Structural and functional MRI data were acquired on a Siemens 3 Tesla PRISMA magnetic resonance imaging scanner equipped with a 64-channel RF head receiver coil. We report details of the imaging sequences following the COBIDAS consortium guidelines (Nichols et al., 2016). Structural T1-weighted images were measured using an MP-RAGE sequence with 176 sagittal slices covering the whole brain, flip angle = 9°, matrix size = 256 × 256 and voxel size = 1 × 1 × 1 mm³. Fieldmaps were acquired using a Siemens gradient echo fieldmap sequence with short echo time (TE) = 5.19 ms and long TE = 7.65 ms (TE difference = 2.46 ms). fMRI data (10 minutes pre-stimulation baseline and 10 minutes concurrent stimulation) were acquired as T2*-weighted gradient echo echo-planar images (EPIs) using a multiband factor of 4, 68 axial slices with an interleaved slice order covering the whole brain including brain stem, repetition time (TR) = 1.4 s, TE = 30 ms, flip angle = 65°, 110 × 110 matrix, field of view = 220 × 220 mm² and voxel size = 2 × 2 × 2 mm³. Additionally, peripheral physiological recordings of the respiratory cycle using the Siemens respiratory belt were planned. Due to the preregistered placement of EGG and ECG electrodes, however, there was not enough space on the body to place the belt without affecting EGG recordings. As the respiratory cycle can also be extracted from electrodes placed on the abdomen (Sayadi, Weiss, Merchant, Puppala, & Armoundas, 2014), we used the EGG measurements to infer the respiratory cycle instead.

#### 2.5.2 Preprocessing

rs-fMRI data was preprocessed using the standardized fMRIPrep pipeline (https://github.com/poldracklab/fmriprep) v20.1.1 (Esteban et al., 2019) [RRID:SCR_016216] based on Nipype (Gorgolewski et al., 2011) [RRID:SCR_002502] and Nilearn (Abraham et al., 2014) [RRID:SCR_001362].

Using this pipeline, each T1-weighted (T1w) volume was corrected for intensity non-uniformity using N4BiasFieldCorrection v2.1.0 (Tustison et al., 2010) and skull-stripped using antsBrainExtraction.sh v2.1.0 (using the OASIS template). Brain surfaces were reconstructed using recon-all from FreeSurfer v6.0.1 (Dale, Fischl, & Sereno, 1999) [RRID:SCR_001847], and the brain mask estimated before was refined with a custom variation of the method to reconcile ANTs-derived and FreeSurfer-derived segmentations of the cortical gray-matter of Mindboggle (Klein et al., 2017) [RRID:SCR_002438]. Spatial normalization to the ICBM 152 Nonlinear Asymmetrical template version 2009c (Fonov, Evans, McKinstry, Almli, & Collins, 2009) [RRID:SCR_008796] was performed through nonlinear registration with the antsRegistration tool of ANTs v2.1.0 (Avants, Epstein, Grossman, & Gee, 2008) [RRID:SCR_004757], using brain-extracted versions of both T1w volume and template. Brain tissue segmentation of cerebrospinal fluid (CSF), white-matter (WM) and gray-matter (GM) was performed on the brain-extracted T1w using FAST (Zhang, Brady, & Smith, 2001) [FSL v5.0.9, RRID:SCR_002823].

Functional data were slice-time corrected using 3dTshift from AFNI v16.2.07 (Cox, 1996) [RRID:SCR_005927] and motion corrected using MCFLIRT (Jenkinson, Bannister, Brady, & Smith, 2002) [FSL v5.0.9]. Distortion correction was performed using fieldmaps processed with FUGUE (Jenkinson, 2003) [FSL v5.0.9]. This was followed by co-registration to the corresponding T1w using boundary-based registration (Greve & Fischl, 2009) with 9 degrees of freedom, using bbregister [FreeSurfer v6.0.1]. Motion correcting transformations, field distortion correcting warp, BOLD-to-T1w transformation and T1w-to-template (MNI) warp were concatenated and applied in a single step using antsApplyTransforms [ANTs v2.1.0] based on Lanczos interpolation.

Physiological noise regressors were extracted by calculating the average signal inside the anatomically-derived CSF and WM masks across time using Nilearn. Framewise displacement (Power et al., 2014) was calculated for each functional run using the implementation of Nipype. Following the recommendation of Power et al. (2014), we calculated the number of volumes per run which exceed a framewise displacement threshold of 0.5 mm. If more than 50% of the total number of volumes exceed this threshold or less than 5 minutes of data below this threshold remain, the respective subject was excluded from further analyses. As our main hypothesis and the accompanying power analysis depend on the imaging data, excluded participants were replaced with new participants until a full set of 40 imaging datasets with two sessions (tVNS and sham) passing quality control was reached.

Spatial smoothing can lead to excessive false positives in smaller brain areas such as the NTS (Frangos et al., 2015; Frangos & Komisaruk, 2017) and increases the risk to mix in signals from outside the respective brain region (Bielczyk, Llera, Buitelaar, Glennon, & Beckmann, 2017). Thus, we initially planned to smooth the whole-brain voxel-based maps with a 6 mm FWHM kernel and keep the unsmoothed data in parallel to extract unsmoothed seed time series from the NTS. However, models to estimate the canonical profiles for stimulation ON- and OFF-cycles based on unsmoothed data showed great instability and did not converge, suggesting that unsmoothed data is not suitable for the modelling approach we had preregistered. Several studies using multivariate pattern analysis have shown that applying spatial smoothing using a small smoothing kernel improves classification accuracy, even though the technique is intended to capitalize on unique spatial information (Gardumi et al., 2016; Mandelkow, de Zwart, & Duyn, 2017). The increase in accuracy is hypothesized to be due to an increase in signal-to-noise ratio (SNR) as reproducible neural contributions to the BOLD signal are spatially autocorrelated and smooth and, thus, can be inferior to noise signal when spatial smoothing is not applied. Importantly, this increase in accuracy also holds true for small regions of interest (Mandelkow et al., 2017). Hence, we smoothed the whole-brain voxel-based maps with a 6 mm FWHM kernel as planned. Due to the small size of our ROI, we also evaluated an NTS time series after smoothing the data with a smaller 4 mm FWHM kernel. Given our voxel size of 2 mm, this was based on recommendations that the minimum FWHM should be twice the size of the voxel (Mikl et al., 2008; Worsley & Friston, 1995). NTS time series extracted from smoothed data showed a Pearson correlation of *r* = .996 indicating negligible differences. Hence, estimating the Time (pre, post) × Stimulation (sham, tVNS) interaction after smoothing with a 6 mm FWHM kernel showed comparable results to the 4 mm FWHM kernel model. In line with our preregistered methods, we also report results from mixed-effects models based on unsmoothed NTS time series.

Respiratory cycle data from the EGG electrodes was preprocessed using BrainVision Analyzer (Brain Products, Germany), FieldTrip (Oostenveld, Fries, Maris, & Schoffelen, 2011) and the PhysIO toolbox (Kasper et al., 2017). In brief, the EGG recordings were read into BrainVision Analyzer and the Scanner Artifact Correction was applied to remove gradient artifacts from the data before submitting exported time series to FieldTrip. As the typical respiratory rate in humans is around 0.3 Hz, the data were downsampled to 50 Hz and bandpass-filtered between 0.1 and 0.6 Hz. The respiratory time series were then read into the PhysIO toolbox and respiratory phase and respiratory volume per time were calculated. By convolution of the respiratory volume per time with the respiration response function (Birn, Smith, Jones, & Bandettini, 2008), the toolbox then generated a nuisance regressor for noise correction. We included this regressor in our MR time series analyses, as respiration influences the activity of the NTS and may affect tVNS-induced brain responses as well (Sclocco et al., 2019).

#### 2.5.3 Statistical analysis

To assess the effect of tVNS on brain dynamics, we investigated the spatiotemporal evolution of brain activation induced by the preset NEMOS stimulation protocol (alternating between ON and OFF every 30 seconds). We then tracked the tVNS-induced activation across the brain using FC measures based on complementary amplitude and frequency information.

##### Hypothesis 1: Temporal evolution of the stimulation effect (main objective)

First, we aimed to model acute effects of tVNS on brain activation based on the three likely temporal activation profiles following stimulation (Figure 3). We reasoned that tVNS either leads to 1) a linear increase in signal amplitude from 0 at t0 s to 1 at t30 s (ramping), 2) an impulse function with a subsequent linear decrease in signal amplitude from 1 at t0 s to 0 at t30 s (decay), or 3) a block-wise shift in signal amplitude to 1 from t0 s to t30 s. In the design matrix, the baseline period preceding the stimulation period was modeled with a constant amplitude of 0. To estimate the stimulation effect on BOLD responses, we generated two design regressors by separately convolving ON and OFF phases of the stimulation (including baseline) with the canonical hemodynamic response function in SPM12. This yielded a set of nine models (Figure 3; three hypothesized design regressors per phase (ON vs. OFF). As we expected tVNS to initially increase brain activation in the NTS, we extracted the mean BOLD time series of the NTS ROI from the acquired rs-fMRI data. To investigate the possible lateralization of the stimulation effect, we extracted two time series from the NTS, one for each hemisphere. We tested if the right and left NTS-based time courses lead to the same winning model. Based on the result of this model comparison, we averaged the NTS time courses over the bilateral ROI mask. Based on a recent publication using a high-resolution MT-weighted sequence at 7T to compile a mask of the NTS (Priovoulos, Poser, Ivanov, Verhey, & Jacobs, 2019), we adapted this mask for usage in functional imaging by enlarging it using the dilation option of FSL Maths [FSL v5.0.9] based on a 9 × 9 × 9 kernel box. However, based on this inflated mask, our statistical models did not converge robustly. Comparing the coordinates provided in Priovoulos et al. (2019) and the coordinates of the provided mask we inflated, we noticed a discrepancy suggesting that the inflated mask mostly captures CSF signal, leading to insufficiently low temporal SNR (tSNR) and poor alignment with the likely position of the NTS in the EPI images across participants. Thus, we constructed a modified mask using WFU PickAtlas (version 3.0.5b), consisting of five box shapes following the coordinates stated in Priovoulos et al. (2019). The first box (dimensions: [3 3 2]) was placed at the MNI coordinates [5 -40 -46], the second box (dimensions: [3 3 2]) placed at the MNI coordinates [5 -41 -49]. The third box (dimensions: [3 3 2]) was placed at the MNI coordinates [3 -42 -52], the fourth box (dimensions: [2 3 2]) placed at the MNI coordinates [2 -43 -55], and the fifth box (dimensions: [2 3 1]) was placed at the MNI coordinates [2 -44 -58]. Importantly, our modified mask more closely captures the shape and coordinates reported by Priovoulos et al. (2019, Figure 2A) and with time series data extracted from our modified mask, the tSNR improved for all 40 participants (Figure 2B, mean tSNR map has been uploaded to NeuroVault for inspection: https://neurovault.org/collections/CHANQVEU/). The notion that low tSNR in the original, inflated NTS mask led to the convergence problems in our models is further supported by running the analyses with time series extracted from a restricted version of the original, inflated mask which only contained the 25% voxels with the highest tSNR. Models based on this restricted version of the inflated mask converged and led to results comparable to those with our modified mask (see Supporting Information - tVNS-induced effects on resting-state fMRI BOLD). We added our modified NTS mask to OSF: https://osf.io/e3gyq/

**Figure 2.**
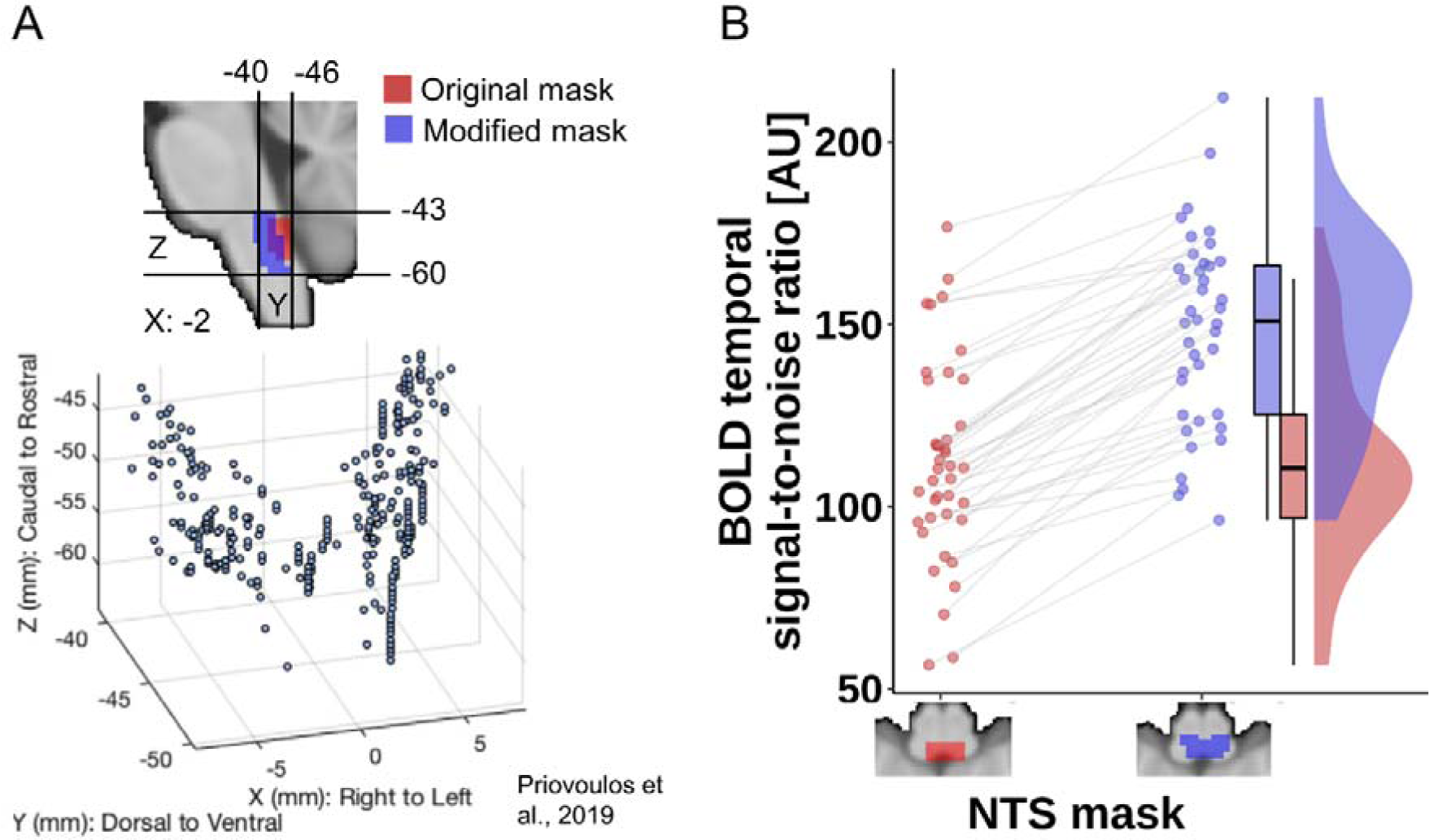
Comparison of the original, inflated NTS mask from Priovoulos et al. (2019, red) and our modified NTS mask (blue) constructed using WFU PickAtlas according to the MNI coordinates reported in Priovoulos et al. (2019). **A:** Comparing the positioning of the new modified NTS mask and the original, inflated mask (upper panel) with the distribution of peak individual voxel intensity along the rostral-caudal axis in MNI space (lower panel) reported by Priovoulos et al. (2019) shows that the modified NTS mask better captures the shape and coordinates (lower panel figure adapted from Priovoulos et al. (2019)). **B:** Temporal signal-to-noise ratio (tSNR) in the modified NTS mask compared to the original, inflated NTS mask is higher for all 40 participants included in the study (mean [95% CI] paired t-test: 36.46 [31.84, 41.08], *t* = 15.96, *p* < .001; the modified NTS mask is available on OSF: https://osf.io/e3gyq/).

**Figure 3.**
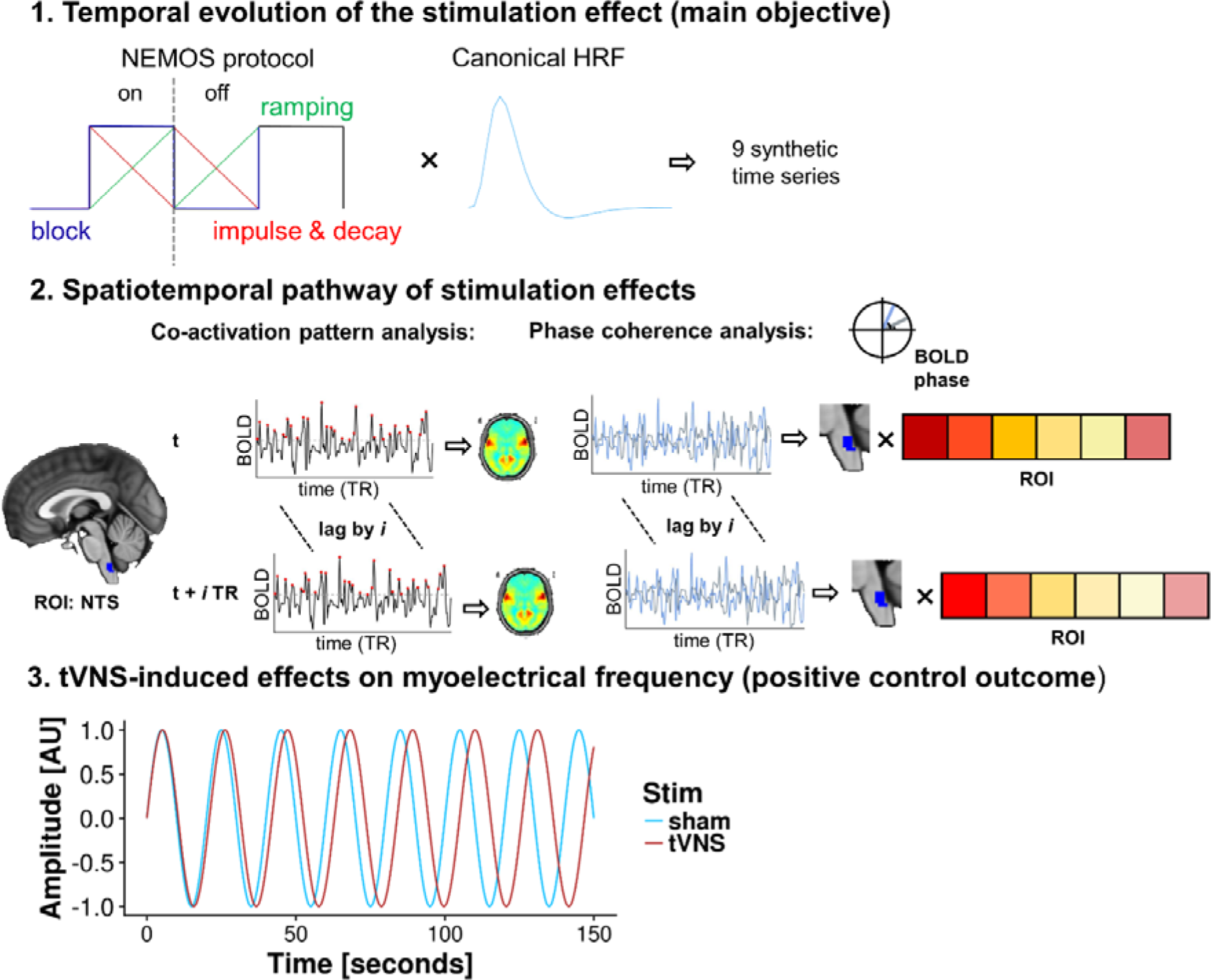
Summary of the key hypotheses and the corresponding analyses. First, we estimated the temporal profile of the tVNS effect for ON and OFF stimulation phases. Second, we mapped the spatiotemporal progression of the tVNS effect using complementary methods that capture functional connectivity. Third, we tested if tVNS induces reliable effects on myoelectrical frequency, which may serve as a positive control of NTS activation via tVNS in future studies.

To identify the best model of tVNS-induced brain activation, time series generated from the convolved design regressors were compared to the extracted time series. As in previous studies (Kroemer et al., 2014, 2016), we used full mixed-effects models, here implemented using the fitlme function in MATLAB, to predict the extracted NTS time series by a set of nine candidate models. Candidate models were composed by combining the potential design regressors. As nuisance regressors, we further included the PhysIO computed respiratory confound regressor, the six realignment parameters, the continuous log-transformed framewise displacement vector, and the WM and CSF regressors as calculated by fMRIPrep. On the participant (second) level of the hierarchy, we modeled random intercepts and slopes for each participant for the ON and OFF stimulation regressors. This enabled us to estimate deviations of each individual from the average group effect (fixed effect). As nuisance regressors, we included the mean framewise displacement^2^ and order of the conditions (tVNS first/sham first; default confound model). The best fitting model over all participants was then identified via model selection based on deviance (i.e., model residuals) of the fMRI time series at first level. Given that all candidate models had the same complexity (i.e., number of free parameters), the best fitting model has the lowest deviance from the NTS time series across the sample. To facilitate comparisons, we calculated the Bayesian information criterion (BIC) with

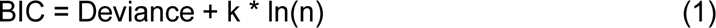

(where k is the number of model parameters and n is the sample size) and the differences in BIC between the best fitting model and other candidates (Kass & Raftery, 1995). A ΔBIC > 10 was considered as strong evidence in favor of the model. If several models fell within this range, this would indicate that the differences between the temporal profiles is negligible. To gain further insights regarding the temporal evolution of the stimulation effect, we additionally inspected and plotted the residuals of the winning model.

Subsequently, we tested an extended confound model as an alternative that might account for additional variance at the participant level. This extended confound model included BMI and sex as these variables might affect the estimation of tVNS effects. We reasoned that the range of age would likely be limited in our sample so that it does not need to be included. Since more nuisance predictors lead to models with higher complexity, we compared the log-likelihood of nested models (default confounds vs. extended confounds) and used the ΔBIC to select the best model for group inferences. tVNS-induced effects were assessed by comparing the model estimates of ON and OFF slopes for tVNS vs. sham sessions. After inspecting the residuals on the participant level, we found a very high correlation between random effects of the slope estimation which suggests that the ON and OFF slopes are statistically inseparable (*r* > .9). Thus, we investigated tVNS-induced effects by contrasting the stimulation and baseline phases of tVNS vs. sham sessions instead.

Lastly, to avoid missing a more appropriate temporal model, we used a “model-free” analysis of the temporal profile of the NTS BOLD time series. We hypothesized that during the tVNS session, the power in higher frequency bands should be increased. We separately estimated the power spectral density of the NTS BOLD time series for each session during baseline (tVNS vs. sham session) and stimulation (tVNS vs. sham session). For statistical analysis, we tested for an interaction between time (pre vs. post stimulation) and stimulation (tVNS vs. sham) on the first level. Again, on the second level of the hierarchy, we modeled random intercepts and slopes for each participant.

##### Hypothesis 2: Spatiotemporal pathway of the tVNS effect

Next, we studied the spatiotemporal dynamics of tVNS-induced brain activation via the NTS across the brain over time. We investigated connectivity between the NTS and other functionally connected ROIs in the brain. To define ROIs, we used an extended version of the Harvard-Oxford atlas (Desikan et al., 2006), which includes the Reinforcement Learning Atlas (https://osf.io/jkzwp/) for extended coverage of subcortical nuclei and the AAL cerebellum ROIs (Tzourio-Mazoyer et al., 2002). Since recent studies have highlighted multiple limitations of sliding windows to estimate dynamics of FC (Hindriks et al., 2016; Preti, Bolton, & Van De Ville, 2017; Tagliazucchi, Siniatchkin, Laufs, & Chialvo, 2016), we used two techniques that allowed us to recover these dynamics without integration over multiple TRs. Critically, these methods are complementary and have been previously shown to recover whole-brain FC dynamics (Deco & Kringelbach, 2016; Di & Biswal, 2015; Kopel, Emmert, Scharnowski, Haller, & Van De Ville, 2017; Liu & Duyn, 2013; Tagliazucchi, Balenzuela, Fraiman, & Chialvo, 2012).

In particular, we used co-activation patterns (CAPs; Liu, Chang, & Duyn (2013), to identify regions that co-activate with the NTS. CAPs are a threshold-based measure of FC between a seed region and the rest of the brain. They reliably track time point-to-time point alterations in “brain states” across participants over a wide range of thresholds (Liu & Duyn, 2013). To extract the CAP of interest, we took the confound-corrected BOLD time series from the NTS seed region (generated within the analysis for Hypothesis 1) during stimulation (tVNS session and sham session) as well as during baseline (tVNS session and sham session). Next, the time points corresponding to the positive peaks of the NTS time series were detected. Time points were included if the amplitude of the signal exceeded a main threshold of the mean amplitude plus two standard deviations (SD), defined for each time series separately. While prior work shows that CAP maps are stable over a wide range of thresholds (Liu & Duyn, 2013), there is no prior work on CAPs extracted from brain stem regions, because they mostly focused on the default mode network. Consequently, it is difficult to judge if CAP maps calculated from a seed in the NTS would fare differently. To test this, we ran an additional sensitivity analysis for the CAP threshold. In this analysis, we tested whether thresholds of SD ±1 result in congruent CAP maps. Therefore, we calculated similarity as well as the Dice and

Jaccard coefficients that can be interpreted based on established guidelines (Fröhner et al., 2019). Lastly, the CAP was obtained by averaging the whole-brain images at these time points. To statistically isolate tVNS effects, we compared the estimated CAPs during baseline (tVNS session vs. sham session), as well as the CAPs during stimulation (tVNS session vs. sham session) on the first level. On the participant (second) level, we modeled random intercepts and slopes for each participant. This enabled us to estimate deviations of each individual from the average group effect (fixed effect). To reveal the cascade of co-activation, we shifted the NTS seed BOLD time series 10 times by one TR and repeated this procedure. This stepwise time-delayed CAP approach allowed us to track the change in spatiotemporal dynamics elicited by the stimulation as compared to sham through the brain.

As tVNS-induced effects may not only alter signal amplitude, complementary methods capitalizing on phase information can help to provide additional insights about the elicited dynamics within the NTS circuit. Thus, we further calculated phase coherence as a measure of FC between the seed region (i.e. NTS) and other brain regions which is independent of the scale of activity (Glerean, Salmi, Lahnakoski, Jääskeläinen, & Sams, 2012). We estimated phase coherence separately for each session during baseline (tVNS vs. sham session) as well as during stimulation (tVNS vs. sham session). To do that, we first normalized the BOLD time series of all regions included in the parcellation built for the initial CAP analysis. Next, to calculate the phase of the signals, we applied the Hilbert transformation. Finally, to obtain the phase coherence values, we used the cosine function on the difference of the phases of each NTS to ROI pair at each TR. This gave us delta FC maps with single TR resolution. To statistically isolate tVNS effects from those of sham, we compared the estimated FC maps during baseline (tVNS session vs. sham session), as well as the FC maps during stimulation (tVNS session vs. sham session) on the first level. On the participant (second) level, we modeled random intercepts and slopes for each participant. This enabled us to estimate deviations of each individual from the average group effect (fixed effect). Similar to the CAP analysis, we then shifted the NTS seed BOLD time series 10 times by one TR and repeated this procedure (Figure 3). This allowed us to identify the exact spatiotemporal signature of the tVNS effect. To control for multiple comparisons in both the CAPs analysis described in the preceding section as well as the phase coherence analysis described here, we used the Benjamini-Hochberg false discovery rate (fdr_bh function with ‘dep’ option as implemented in MATLAB R2017a).

Additionally, we counted the number of peaks which are determined after thresholding the signal using the same procedure as described above (calculation of CAPs) in each of the four conditions. For statistical analysis, we used the estimated number of peaks during baseline (tVNS session vs. sham session), as well as the frequencies during stimulation (tVNS session vs. sham session) on the first level. For the sparse confound model, we added order of stimulation conditions (tVNS first/sham first) as a nuisance regressor at the second level. We hypothesized that the number of peaks should be highest in the tVNS condition (time × stimulation interaction).

### 2.6 Electrogastrogram

The EGG non-invasively measures electrical activity reflecting the rhythmic contractions of the stomach. This gastric rhythm is initiated by pacemaker cells seated in the mid-to-upper corpus of the stomach (O’Grady et al., 2010). These pacemaker cells receive efferent vagal inputs from the NTS via the dorsal motor nucleus of the vagus (Koch & Stern, 2004). Then, pacemaker currents propagate to the rest of the stomach, ensuring that muscle cells in the stomach contract in an orchestrated manner for digestion (Rebollo et al., 2018).

#### 2.6.1 Data acquisition

EGG data were acquired according to the protocol described by Rebollo et al. (2018). Briefly, we recorded EGG and associated triggers with BrainVision Recorder (Brain Products, Germany) using four bipolar standard adhesive electrocardiogram (ECG) electrode pairs (eight electrodes in total) connected to a BrainAmp amplifier (Brain Products, Germany). For EGG acquisition, the electrodes were placed in a bipolar montage (four pairs) in three rows over the abdomen to keep MR gradient artifacts at a minimum. EGG was acquired at a sampling rate of 5000 Hz with a low-pass filter of 1000 Hz and no high-pass filter (DC recordings). To evaluate efferent effects of tVNS, we continuously recorded EGG inside the scanner including baseline and stimulation periods. To mark the beginning and end of the baseline as well as the stimulation phase, triggers were sent using Psychtoolbox (http://psychtoolbox.org/).

In addition to EGG, we also measured ECG using three bipolar electrodes set on both sides below the clavicula as well as on the left side below the ribcage. The ECG is no measure of interest for the current report.

#### 2.6.2 Preprocessing

For preprocessing of the EGG data, we employed the analysis scripts written by Rebollo et al. (2018), which are available on GitHub (https://github.com/irebollo/stomach_brain_Scripts). The preprocessing was based on the FieldTrip toolbox (Oostenveld, Fries, Maris, & Schoffelen, 2011) implemented in MATLAB. Accordingly, data were low-pass filtered with a cut-off frequency of 5 Hz and downsampled to 10 Hz. Given that the gastric frequency is known to be ∼0.05 Hz and MR gradient artifacts only affect signals >10 Hz, there was no specific artifact correction procedure needed for gradient artifacts (Rebollo et al., 2018). Next, to identify the individual EGG peak frequency, a fast fourier transform was run for each EGG channel separately to calculate the spectral density of the signal between 0 Hz and 0.1 Hz using MTMFFT as implemented in FieldTrip. The frequency range of interest was 0.033–0.066 Hz and EGG peak identification was based on two criteria:

The respective peak had to have a power > 15 μV^2^. If peak estimates in more than one EGG channel exceeded this measure, the channel containing the sharpest peak was selected. After channel selection, the respective channel data were bandpass-filtered to isolate the individual peak gastric frequency using a MATLAB-based FIR filter centered at the respective EGG peaking frequency with a filter width of ±0.015 Hz and a filter order of 5. EGG data were excluded from further analysis if the spectral power in the frequency range of interest did not exceed 15 μV^2^ in any of the EGG channels or no sharp frequency peak could be identified from any of the EGG channels. Excluded EGG sessions were not replaced with new participants as EGG data were not related to the main hypothesis, but of relevance for the positive outcome control only. To maximize power, we instead included single sessions that passed quality control in group analyses using mixed-effects analysis.

#### 2.6.3 Statistical analysis

##### Hypothesis 3: tVNS-induced effects on myoelectrical frequency (positive control outcome)

For statistical analysis, we compared the pre/post (baseline/stimulation) peak frequencies of the tVNS and sham measurements. As the impact of possible confound variables on EGG is not clear to date, we tested two mixed-effects models with different model complexity. Again, we calculated the deviance and BIC at the group level as described in 2.5.3. (Statistical analysis, temporal evolution of the tVNS effect).

For EGG group analysis, we used the estimated frequencies during baseline (tVNS session vs. sham session), as well as the frequencies during stimulation (tVNS session vs. sham session) on the first level. As for the other hypotheses, we compared the extended confound model to the default confound model. Finally, after the winning model was identified, the interaction effect of Time (pre/post) × Stimulation (tVNS/sham) was evaluated to test for the hypothesized slowing in gastric frequency after tVNS.

## 3. Results

### 3.1 Temporal evolution of the stimulation effect

To estimate the effect of tVNS on BOLD responses in the NTS, we used full mixed-effects models. We predicted NTS time series extracted from smoothed rs-fMRI data by a set of nine candidate models, tested for a possible lateralization of the stimulation effect, and identified the best confound model.

#### Model comparisons between candidate models for the temporal profile of the tVNS effect

To compare the model fit between our 9 candidate models, we calculated the BIC for each model and the ΔBIC between the best fitting model and all other candidate models. First, we compared models based on time series extracted from the right and left NTS, separately. For both sides, predictions based on the candidate model “block ON/block OFF” had the lowest deviance from the extracted data (deviances for all models based on the left, right and bilateral NTS can be found in Supplementary Table S1). Thus, we averaged the time series from the left and right NTS and following analyses were based on the bilateral NTS ROI.

Based on the time series extracted from the bilateral NTS ROI, we found strong evidence in favor of the block ON/block OFF model (ΔBIClowest = 992 > 10 for the comparison between the block ON/block OFF and the ramping ON/block OFF model; ΔBIChighest = 2085 for the comparison between the block ON/block OFF and the decay ON/ramping OFF model; Figure 4A). To evaluate confounds at the participant level, we compared the model fit between the default and the extended confound model and found strong evidence in favor of keeping the sparse model (ΔBIC = 20.97 > 10). In contrast, using unsmoothed time series extracted from the bilateral NTS ROI, models did not converge and corresponding model comparisons were not interpretable.

**Figure 4.**
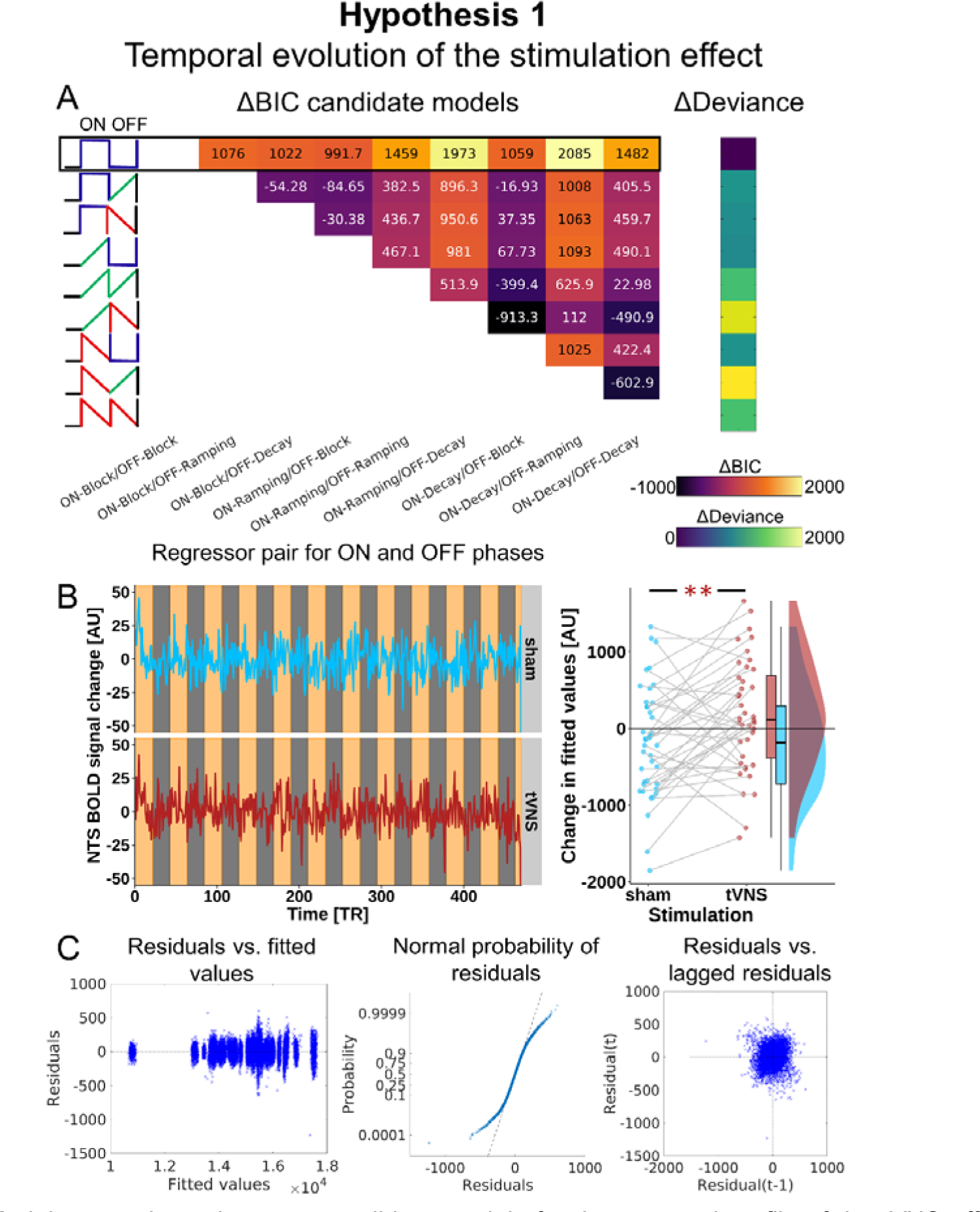
Model comparisons between candidate models for the temporal profile of the tVNS effect. **A:** We calculated delta deviance (compared to the winning model) and delta Bayesian Information Criterion (BIC) for mixed-effects models predicting NTS time series based on the stimulation ON and OFF pairs of design regressors (depicted on the x- and y-axes). The model based on the block ON/block OFF regressor pair (first row, black frame) had the lowest deviance and highest delta BIC with every other possible regressor combination (BIC_lowest_ = 992 > 10). **B:** Mean NTS BOLD signal change (left plot) for sham stimulation (blue, uppe^Δ^r panel) and tVNS (red, lower panel). Light orange areas mark periods recorded during ON phases (either tVNS or sham stimulation, respectively). Light grey areas mark OFF phases. The block ON/block OFF regressor pair best describes the temporal profile of the time series, but coefficients for the block ON/block OFF regressor pair showed a correlation of r = .99, indicating that ON and OFF phases cannot be statistically separated. Comparing stimulation and baseline phases for tVNS vs. sham sessions instead (right plot), shows an increase in resting-state BOLD signal in the NTS following tVNS compared to sham stimulation. The y-axis depicts the demeaned, fitted values for the Time (pre, post) × Stimulation (sham, tVNS) interaction (*t* = 2.97, *p*_Satterthwaite_ = .005). **C:** Residual plots assessing residuals versus fitted values, normality, and autocorrelation of the residuals (from left to right) for the winning model. Whereas the residuals do not correlate with their own lagged values, the normality plot shows heavy tails indicating the presence of many very negative or very positive residual values as further indicated by the clusters on the far left and far right in the residuals versus fitted values plot.

#### tVNS-induced effects on resting-state fMRI BOLD

We did not find a significant block ON × Stimulation (sham, tVNS) interaction (Figure 4B left; FWHM 4mm smoothed data: *p*_Satterthwaite_ = .46, unsmoothed data: *p*_Satterthwaite_ = .26) or block OFF × Stimulation interaction (FWHM 4mm smoothed data: *p*_Satterthwaite_ = .54; unsmoothed data: *p*_Satterthwaite_ = .18). However, in conflict with our preregistered analysis, models did not converge (outputs are provided on OSF: https://osf.io/zmau8/). Therefore, we inspected the model residuals and covariance matrices and found a correlation between the block ON and block OFF random slopes (*r* = .99) that precluded robust separation. Thus, we investigated the tVNS-induced effects by comparing the full stimulation and baseline phases by predicting the bilateral NTS time series based on the Time (pre, post) × Stimulation (sham, tVNS) interaction and the sparse confound model. We found that tVNS compared to sham led to a significant increase in BOLD signal change (Figure 4B right; mean Time × Stimulation: 179.44 [57.15, 301.72], *t* = 2.97, *p*_Satterthwaite_ = .005) indicating that activity within the bilateral NTS increases for the stimulation phase compared to the baseline phase when applying tVNS versus sham stimulation. The empirical effect size was *d_z_* = 0.47 [0.12, 0.84] (correlation between tVNS and sham sessions: *r* = .45 [.18, .68]) with the CI including the recovered *d_z_* = 0.79 as well as the correlation of *r* = .57 between sessions that our simulations were based on. Using data smoothed with a 6 mm FWHM kernel, we obtained comparable results (mean Time × Stimulation: 161.45 [19.19, 303.71], *t* = 2.30, *p*_Satterthwaite_ = .027). Using unsmoothed data, the model failed to converge and the Time × Stimulation interaction was not significant (*t* = 1.0, *p*_Satterthwaite_ = .3).

#### Model-free analysis of the temporal profile of the NTS BOLD time series

To test for an increase in power of NTS BOLD time series for tVNS versus sham stimulation, we estimated the power spectral density (dB) of the confound-corrected NTS BOLD time series. We used a linear mixed-effects model predicting power of NTS BOLD based on the interaction Time (pre, post) × Stimulation (sham, tVNS) over frequency bins ranging from .01 to .15 Hz. We found no significant effect of tVNS on NTS BOLD power (Supplementary Figure S2; *t* = 0.10; *p*_Satterthwaite_ = .9). Similarly, we found no significant effect of tVNS on NTS BOLD power based on unsmoothed NTS time series (*t* = -0.25, *p*_Satterthwaite_ = .8).

### 3.2 Spatiotemporal pathway of stimulation effects

To unravel the pathway of signaling associated with the stimulation, we tested for regions whose activity is associated with the activity within the NTS, statically as well as in a time-lagged manner. We tested for tVNS effects based on signal amplitude by performing incremental calculations of CAPs with increasing lag. Furthermore, we calculated static and dynamic phase-coherence to determine changes in frequency that are orthogonal to changes in amplitude.

#### tVNS-induced effects on co-activation patterns

First, we tested for regions which show a significant static connectivity with the NTS across all sessions (sham, tVNS) and phases (pre, post) by averaging the estimated CAP maps and calculating a one sample t-test for each ROI. After multiple comparison correction, 149 regions showed a significant static connectivity with the NTS (Figure 5; the only regions which did not show a significant static connectivity with the NTS were the subcallosal gyrus, right cerebellar culmen, and the nodule of the cerebellar vermis).

**Figure 5.**
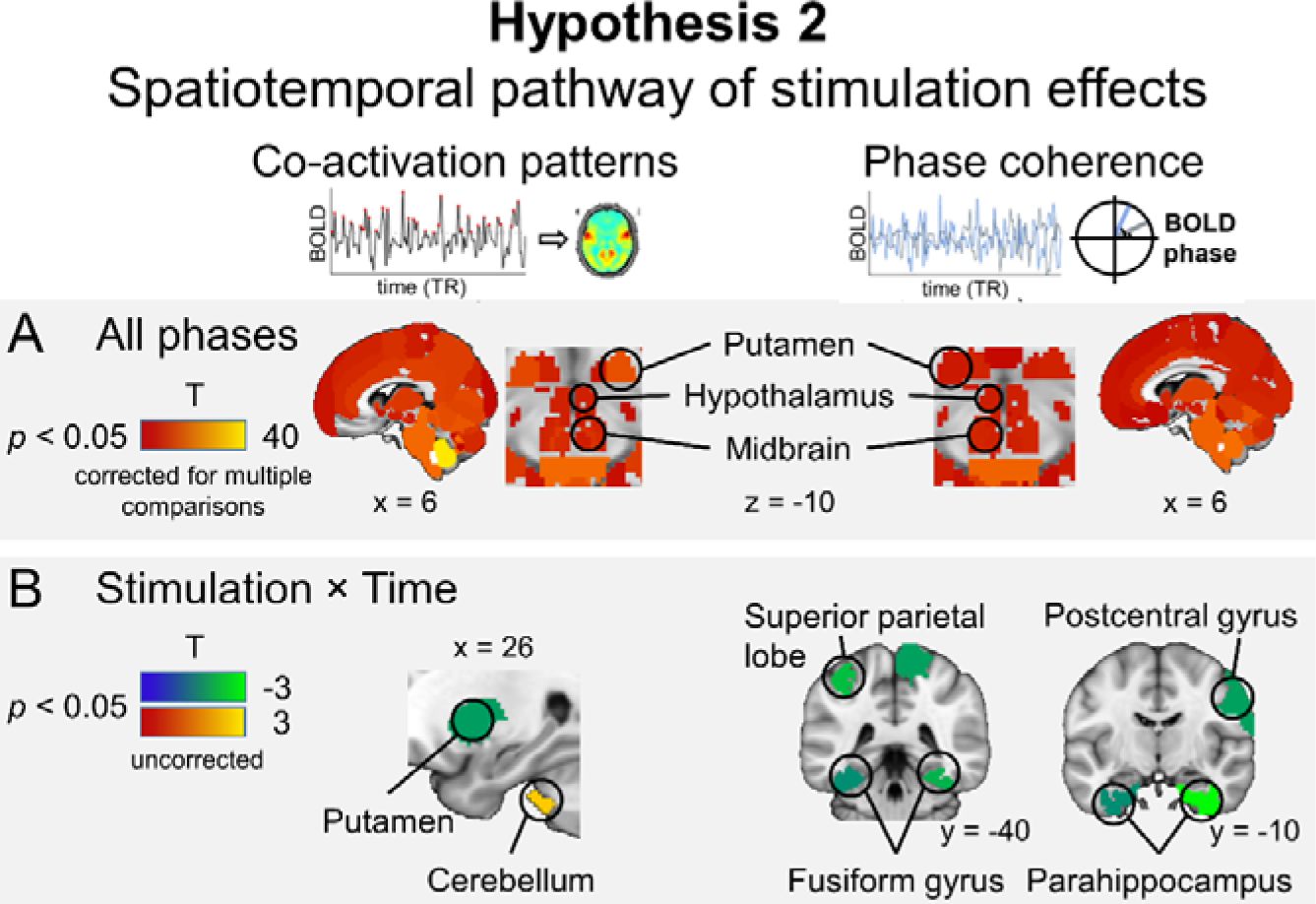
Regions showing static functional connectivity (FC) with the NTS based on co-activation patterns (CAPs, left side) and phase coherence (right side). **A:** An extended network of 149 (CAPs) and 145 (phase coherence) regions, respectively, shows static FC with the NTS (one-sample t-test; all regions *p*_FDR-corrected_ < .05) across all phases (baseline and stimulation in both sessions). **B:** No regions showed altered FC after correction for multiple comparisons. However, tVNS modulated CAPs in the putamen and cerebellum, and phase coherence in the right postcentral gyrus, left superior parietal lobe, bilateral parahippocampal gyrus, bilateral fusiform gyrus, and the left lateral occipital cortex (Stimulation × Time interaction; all regions *p*_uncorrected_ < .05).

To test for the tVNS effect on CAPs, we used mixed-effects models predicting the estimated CAP based on the Time (pre, post) × Stimulation (sham, tVNS) interaction for each ROI in the extended Harvard-Oxford atlas and each time shift. Contrary to our hypothesis, we found no significant effects of tVNS on static or dynamic connectivity based on CAPs after correction for multiple comparisons (supplementary Figure S3; mean CAP maps have been uploaded to NeuroVault for inspection: https://neurovault.org/collections/CHANQVEU/). Without correcting for multiple comparisons, we found a significant Time × Stimulation interaction in the right putamen (*t* = -2.23) and the right cerebellar tonsil (*t* = 2.54; Figure 5).

#### Sensitivity analysis for the co-activation pattern threshold

To test whether different SD thresholds lead to congruent CAP maps, we calculated similarity, Dice and Jaccard coefficients for CAP maps generated based on thresholds of SD = 2 ± 1. We found an excellent overlap (Dice coefficient: SD1/SD2 = .98, SD1/SD3 = .96, SD2/SD3 = .96; Jaccard coefficient: SD1/SD2 = .96, SD1/SD3 = .93, SD2/SD3 = .93) and similarity (similarity: SD1/SD2 = .99, SD1/SD3 = .91, SD2/SD3 = .94), respectively, between the CAP maps resulting from different SD thresholds, indicating that our results are not dependent on the SD threshold.

#### tVNS-induced effects on phase coherence

First, we tested for regions which show a significant phase coherence with the NTS across all sessions and phases by averaging the estimated phase coherence maps and calculating a one sample *t*-test for each ROI. After multiple comparison correction, 145 regions showed a significant phase coherence with the NTS (Figure 5; the only regions which did not show a significant phase coherence with the NTS were the bilateral posterior part of the superior temporal gyrus, left middle temporal gyrus, bilateral inferior temporal gyrus, right fusiform cortex and the cerebellar culmen).

To test for the tVNS effect on phase coherence, we used mixed-effects models predicting the estimated phase coherence based on the Time × Stimulation interaction for each ROI in the extended Harvard-Oxford atlas and each time shift. Analogous to the CAP-based analysis and, again, contrary to our hypothesis, we found no significant effects of tVNS on static and dynamic connectivity based on phase coherence (supplementary Figure S4; mean phase coherence maps have been uploaded to NeuroVault for inspection: https://neurovault.org/collections/CHANQVEU/). Without correcting for multiple comparisons, we found a significant Time × Stimulation interaction in the right postcentral gyrus (*t* = -2.26), left superior parietal lobe (*t* = -2.49), left lateral occipital cortex (*t* = -2.11), bilateral parahippocampal gyrus (right: *t* = -2.96, left: *t* = -2.11), and the bilateral fusiform cortex (right anterior temporal fusiform cortex: *t* = -3.19, right posterior temporal fusiform cortex: *t* = -2.95, right temporal occipital fusiform cortex: *t* = -2.40, left posterior temporal fusiform cortex: *t* = -2.10; Figure 5).

#### tVNS-induced effects on the number of peaks determining the co-activation patterns

To test for the tVNS effect on the number of peaks determined after thresholding the NTS signal for the CAP calculation, we used a mixed-effects model predicting the estimated number of peaks based on the Time × Stimulation interaction. Again, we found no significant effect of tVNS on the number of peaks after thresholding the NTS signal (Supplementary Figure S5; *M* Δ_sham_ = 1.3, *M* Δ_tVNS_ = 0.55, *t* = -0.77, *p*_Satterthwaite_ = .44).

### 3.3 tVNS-induced effects on gastric myoelectrical frequency

To investigate whether gastric myoelectrical frequency is modulated by tVNS, we used a mixed-effects model predicting gastric frequency based on the Time × Stimulation interaction. After initial quality control, data from two participants had to be excluded as the spectral power in the frequency range of interest did not exceed the preregistered threshold of 15μV^2^. We further had to exclude single sessions from 14 participants (*N*_sham_ = 7, *N*_tVNS_ = 7) for whom no sharp frequency peak could be identified. This yielded the final sample of N = 38 participants with at least one session that passed quality control (N = 24 participants with data from both sessions).

First, we compared the model fit between the default and the extended confound models by calculating deviance and ΔBIC between both models. We found evidence in favor of keeping the sparser default confound model (ΔBIC = 5.03).

Contrary to our hypothesis, we found no significant effect of tVNS on gastric myoelectrical frequency (Figure 6; *t* = 0.67, *p*_Satterthwaite_ = .51).

**Figure 6.**
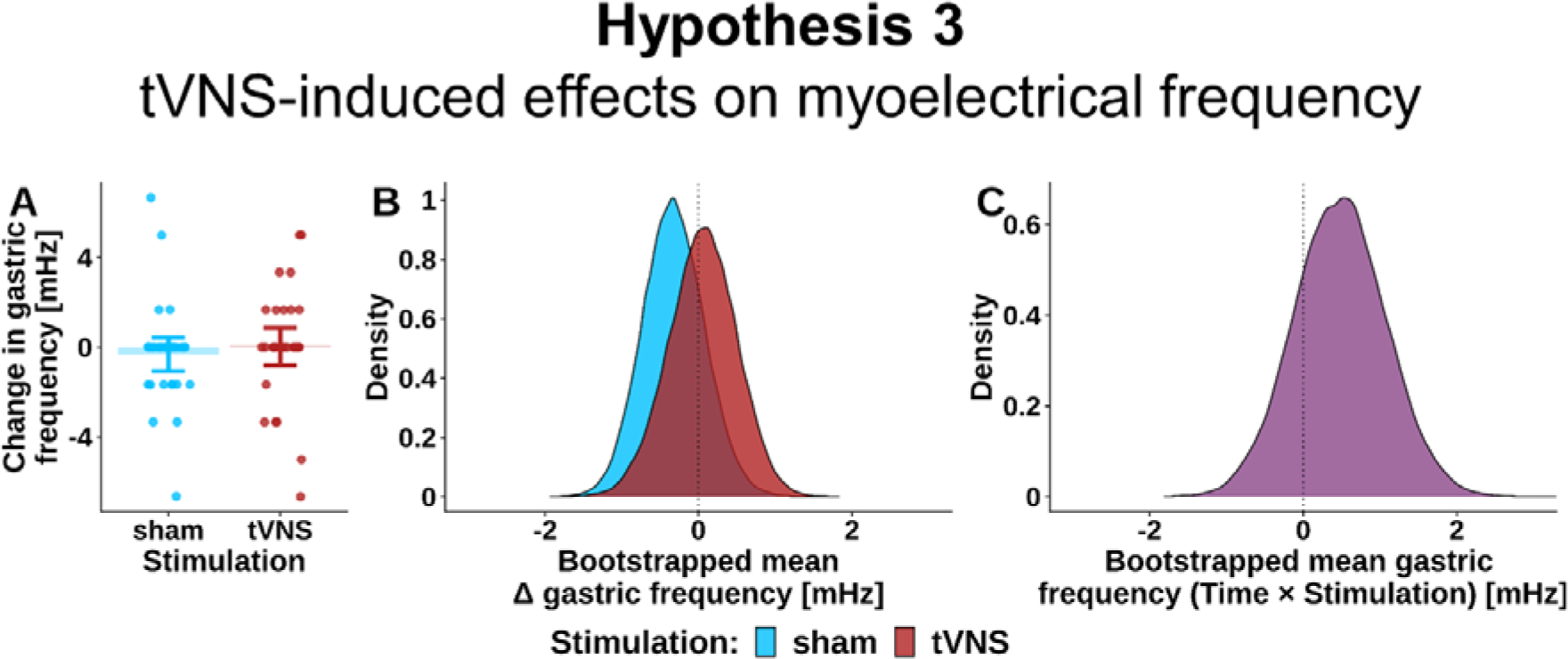
No effect of tVNS compared to sham stimulation on gastric myoelectric frequency. **A:** Change in gastric frequency (post - pre). **B:** Bootstrapped delta gastric frequency distributions (post - pre) for tVNS (red) and sham stimulation (blue) for all data points included in the analysis (including single sessions from 14 participants). **C:** Interaction between Time (pre, post) and Stimulation (sham, tVNS) for all participants with data from two sessions. In contrast to our hypothesis and a previous study from our group (Teckentrup et al., 2020), we found that gastric frequency was not decreased following tVNS compared to sham stimulation.

## 4. Discussion

tVNS is a promising non-invasive brain stimulation technique, but elicited dynamics in the brain are still largely elusive. Here, we applied tVNS versus sham stimulation in 40 healthy participants using a single-blind randomized cross-over design concurrently to resting-state fMRI and EGG recordings. Model comparisons showed that from a set of 9 candidate models, a block-wise shift in signal amplitude after stimulation fitted the NTS time series best. Notably, random effects of ON and OFF stimulation slopes showed a high correlation, indicating that these phases cannot be reliably separated with fMRI BOLD. In line with our first hypothesis, tVNS compared to sham stimulation increased activation in the NTS. Nevertheless, these results were contingent on smoothing and accurate placement of the mask to avoid signal dropout. Contrary to our second hypothesis, we found no evidence that the tVNS-induced changes in activation propagate downstream as measured by CAPs and phase coherence. Also, contrary to our third hypothesis, we found no evidence for a decrease in gastric myoelectrical frequency after tVNS. Thus, acute tVNS can perturb brain signaling in the NTS, but further research using modified protocols is necessary to evaluate downstream effects of the stimulation as well as efferent effects on gastric motility.

Confirming our first hypothesis, we found an increased activity in the bilateral NTS during tVNS compared to sham stimulation. As suggested by our model comparison of candidate temporal profiles, the stimulation likely induces a general shift in NTS activity. Corroborating this result, the confidence interval for the empirical effect size of the stimulation effect included the effect size recovered from our initial simulation. The result fits well with previous studies investigating effects of tVNS on brain activation which mostly showed bilateral (Yakunina et al., 2017; Sclocco et al., 2019) or ipsilateral increases of activation in the NTS (Frangos et al., 2015; Frangos & Komisaruk, 2017). The NTS is the recipient of the majority of vagal afferent projections in the brain stem. Consequently, successful tVNS should modulate signaling within this region. Still, several other studies either found a decrease or no change in activation in the NTS (Dietrich et al., 2008; Kraus et al., 2013; Frangos & Komisaruk, 2017; Badran et al., 2018). This might be explained by the site of stimulation, as stimulating the cymba conchae has been shown to elicit NTS activation most reliably (Yakunina et al., 2017), but these studies either applied tVNS to the tragus or the neck. Furthermore, it has been argued that studies showing increases in NTS activation had optimized their power and sensitivity to observe changes in NTS activity by tuning acquisition parameters (Sclocco et al., 2019) or statistical analyses (Fang et al., 2017). Using our preregistered NTS mask which we inflated for usage with fMRI BOLD data, we noticed a substantial signal loss in the preprocessed rs-fMRI data. Thus, we generated a mask based on the coordinates reported by Priovoulos et al. (2019) with better coverage of gray matter, which improved coverage of the NTS in the normalized EPI data and significantly increased tSNR in all participants (total increase in tSNR of 32.5%). As a small ROI in the brain stem, the NTS is still a challenging region to measure robustly with fMRI due to inherently low SNR (Beissner, 2015; Sclocco, Beissner, Bianciardi, Polimeni, & Napadow, 2018) and cardiorespiratory noise (Brooks, Faull, Pattinson, & Jenkinson, 2013). Accordingly, even when controlling for physiological noise, analyzing our model residuals uncovered heavy tails in the distribution of the data (Kasper et al., 2017). To avoid undue influence of heavy tails, data trimming is a common approach (Wilcox, 2005). Future studies modeling stimulation-induced changes in brain signaling may therefore evaluate trimming as a means to improve robustness. Still, with these issues being addressed in an unbiased manner, our results confirmed the hypothesized increase in NTS activation after tVNS.

In contrast to our second hypothesis, we did not find evidence for effects of tVNS on connectivity between downstream brain regions and the NTS, neither based on signal amplitude using CAPs, nor based on frequency using phase coherence. So far, only few studies investigated connectivity changes after tVNS and typically focused on patient groups with depression (Fang et al., 2017; Liu et al., 2016; Tu et al., 2018; Wang et al., 2018) and migraine (Garcia et al., 2017) after chronic treatment with tVNS. Analyzing static FC across all phases showed that both CAPs and phase coherence captured an extensive network associated with the NTS. However, tVNS did not robustly modulate this NTS FC network (i.e., no significant effects after multiple comparison correction). Individual variability in tVNS-induced effects might have contributed to variability in downstream connectivity effects and future studies on connectivity changes will likely need to recruit larger samples to resolve potentially low to moderate effects on NTS-based FC. Collectively, our results do not provide support for large effects on NTS-based FC at the group level.

In contrast to our third hypothesis, we also did not find evidence for effects of tVNS on gastric myoelectric frequency. This finding is inconsistent with previous studies from our group (Teckentrup et al., 2020) as well as others (Hong et al., 2018). Several differences in the study protocol might have contributed to the divergence from previous work. In the current study, we assessed changes in gastric frequency during 10 minutes of stimulation, whereas we had observed changes in frequency during 30 minutes of stimulation in our previous study. Thus, changes in gastric frequency as measured via EGG might only occur after a temporal delay. However, also Hong et al. (2018) had stimulated for only 10 minutes and found a decrease in gastric frequency after tVNS. Crucially, they measured gastric motility invasively in the pylorus and corpus of the stomach with stronger effects in the pylorus region and non-invasive EGG might reflect different sources, leading to attenuated effects (Wolpert, Rebollo, & Tallon-Baudry, 2020). Moreover, both previous studies stimulated the left branch of the vagus nerve while we stimulated the right branch in the current study. Recent findings in animals showed that the right branch projects to the dopaminergic midbrain (Han et al., 2018) and the differential innervation of mechano- and chemoreceptors in the gut by the left and right vagus nerve endings lends additional credibility to side-specific effects of tVNS (Wang, de Lartigue, & Page, 2020). Thus, changes in gastric frequency might be boosted after left-sided compared to right-sided tVNS. Furthermore, the metabolic state of participants might have played a role as participants in our previous study took part after >4h of fasting, whereas participants in the current study were neither hungry nor full. Importantly, the absence of changes in gastric frequency after right-sided tVNS does not preclude changes in coupling between stomach and brain signaling (Rebollo et al., 2018), which might show instant changes. Taken together, our hypothesis of a decrease in gastric frequency instantly after tVNS was not confirmed, and further studies will need to investigate possible effects of time delay, metabolic state, and lateralization on an alleged biomarker of the stimulation.

Our study has several strengths and limitations which should be addressed in future work. Based on an a priori power analysis and simulations of the expected stimulation effects, we ran a well-powered study including a state-of-the-art sham stimulation condition as control (Farmer et al., 2020) and comprehensive measures to restrict and correct for possible sources of (physiological) noise. We further report an effect size for tVNS-induced effects on brain signaling in the NTS to inform future hypothesis-driven research. Still, tVNS-induced effects were dependent on smoothing and exact placement of the mask. Applying spatial smoothing has been shown to be beneficial in multivariate pattern analysis (Gardumi et al., 2016; Mandelkow et al., 2017) which has been attributed to an increase in SNR (Mandelkow et al., 2017). We showed that using our modified NTS mask improved tSNR in all participants. Compared to the original mask, our modified mask is still based on the MNI coordinates reported by Priovoulos et al. (2019), yet aligns better with the processed EPI data provided by fmriprep after normalization. Importantly, a restricted version of the original, inflated mask which only contained the 25% voxels with the highest tSNR led to results comparable to those with the modified mask. This highlights the importance of using accurately placed masks for time series modeling in small brain regions that are close to non-brain tissue as our preregistered mixed-effects models only converged when tSNR was sufficiently high. Still, the tVNS effects based on the modified NTS mask should be validated in further studies using concurrent stimulation and fMRI. To address the differences in results on changes in dynamic FC after tVNS, more studies with large sample sizes are needed. Based on our results, future studies using a within-participant design should preferably include more than 40 participants to identify robust changes after stimulation, specifically in functional connectivity.

To summarize, tVNS is a promising technique to non-invasively alter brain function. Here, in line with most previous reports, we showed that tVNS increases fMRI BOLD activation in the NTS. Extending prior and informing future work using tVNS, we found that NTS BOLD activation during stimulation is best characterized by a simple block model describing a general increase in activity during tVNS. Our results further provide substantial evidence for using bilateral NTS masks as we found no evidence supporting a lateralization of stimulation effects in the NTS with fMRI. However, our study could not identify a more fine-grained spatiotemporal profile of the stimulation as intended because tVNS-induced changes did not exceed stringent correction levels. Additionally, we highlight that tVNS effects are sensitive to spatial smoothing and placement of the mask to avoid insufficient tSNR for statistical inference. Likewise, we found no evidence for a robust decrease in gastric frequency during tVNS. While this observation might be due to various differences to previous studies (e.g., the metabolic state of the participant, side of the stimulation, or a delay in onset of efferent effects on the stomach), it indicates that the use of the EGG as a biomarker of tVNS requires further validation. Collectively, our results support the notion that tVNS is a promising technique to perturb brain signaling in a well-defined neuroanatomical circuit that is hard to target with alternative methods. Thus, our work serves as a comprehensive reference to advance future research on brain dynamics after tVNS.

## Acknowledgements

We thank Sophie Müller, Vinzent Wolf, Franziska Müller, Wy Ming Lin, Franziska Kräutlein and Corinna Schulz for help with data acquisition. The study was supported by the University of Tübingen, Faculty of Medicine fortüne grant #2453-0-0. MPN received salary support from the Else Kröner-Fresenius-Stiftung, grant #2017-A67 and the University of Tübingen, Faculty of Medicine ‘forschungsorientierte Gleichstellungsförderung’ #2605-0-0 awarded to NBK. NBK received additional support from the Daimler and Benz Foundation, grant 32-04/19. HJ was supported by the University of Tübingen, Faculty of Medicine fortüne grant #2487-1-0.

## Author contributions

NBK was responsible for the concept and design of the study. VT & RH performed pilot measurements to evaluate usage of the stimulator within the scanner. VT & SN collected data. NBK, VT, MK & HJ conceived the method and VT & NBK processed the data. VT performed the data analysis and NBK, MK & HJ contributed to analyses. VT, NBK, SN & MPN wrote the Stage 1 manuscript and VT & NBK wrote the Stage 2 manuscript. All authors contributed to the interpretation of findings, provided critical revision of the manuscript for important intellectual content and approved the final version for publication.

## Data and Code availability statement

The code used to calculate the power analysis described in section 2.2 is available for download from https://osf.io/v8ngb/. The codes generated for planned analyses in the study (https://osf.io/v8ngb/) and summary statistics of the planned analyses (https://osf.io/zmau8/) are uploaded to the same repository. In accordance with the grant used to fund the study (Tübingen’s fortune #2453-0-0), the acquired raw (nifti) data will be made openly available via a public repository one year after the completion of the study.

## Footnotes

1 Scripts used to run the simulations described in this section can be downloaded here: https://osf.io/v8ngb/

2 see https://neurostars.org/t/confounds-from-fmriprep-which-one-would-you-use-for-glm/326

