## Supporting Information for "Brain signaling dynamics after vagus nerve stimulation"

*tVNS-induced effects on resting-state fMRI BOLD (restricted version of original NTS mask)*

We investigated if low tSNR in the original, inflated NTS mask led to convergence problems in our models by running the analysis (Time (pre, post) × Stimulation (sham, tVNS)) with time series extracted from a restricted version of the original, inflated mask which only contained the 25% voxels with the highest tSNR. Using this restricted mask, the results were comparable to the analysis based on our new NTS mask: mean [95% CI] Time × Stimulation: 221.67 [114.81, 328.54], *t* = 4.20, *p*_Satterthwaite_ < .001. This corroborates the notion that low tSNR led to the convergence problems using time series extracted from the original NTS mask.

**Table S1.** Deviance values for models predicting time series extracted from the NTS right, left and bilateral (modified NTS mask) based on all temporal profile regressor pairs. The block ON/block OFF model shows the lowest deviance values for all three masks. Thus, in subsequent analyses, time series from the left and right NTS were averaged.

| ON OFF | block block | block ramping | block decay | ramping block | ramping ramping | ramping decay | decay block | decay ramping | decay decay |
| --- | --- | --- | --- | --- | --- | --- | --- | --- | --- |
| NTS right | 428132 | 429078 | 429031 | 429007 | 429423 | 429879 | 429061 | 429966 | 429431 |
| NTS left | 438746 | 439465 | 439436 | 439408 | 439729 | 440086 | 439444 | 440137 | 439731 |
| NTS bilateral | 426293 | 427369 | 427315 | 427285 | 427752 | 428266 | 427352 | 428378 | 427775 |

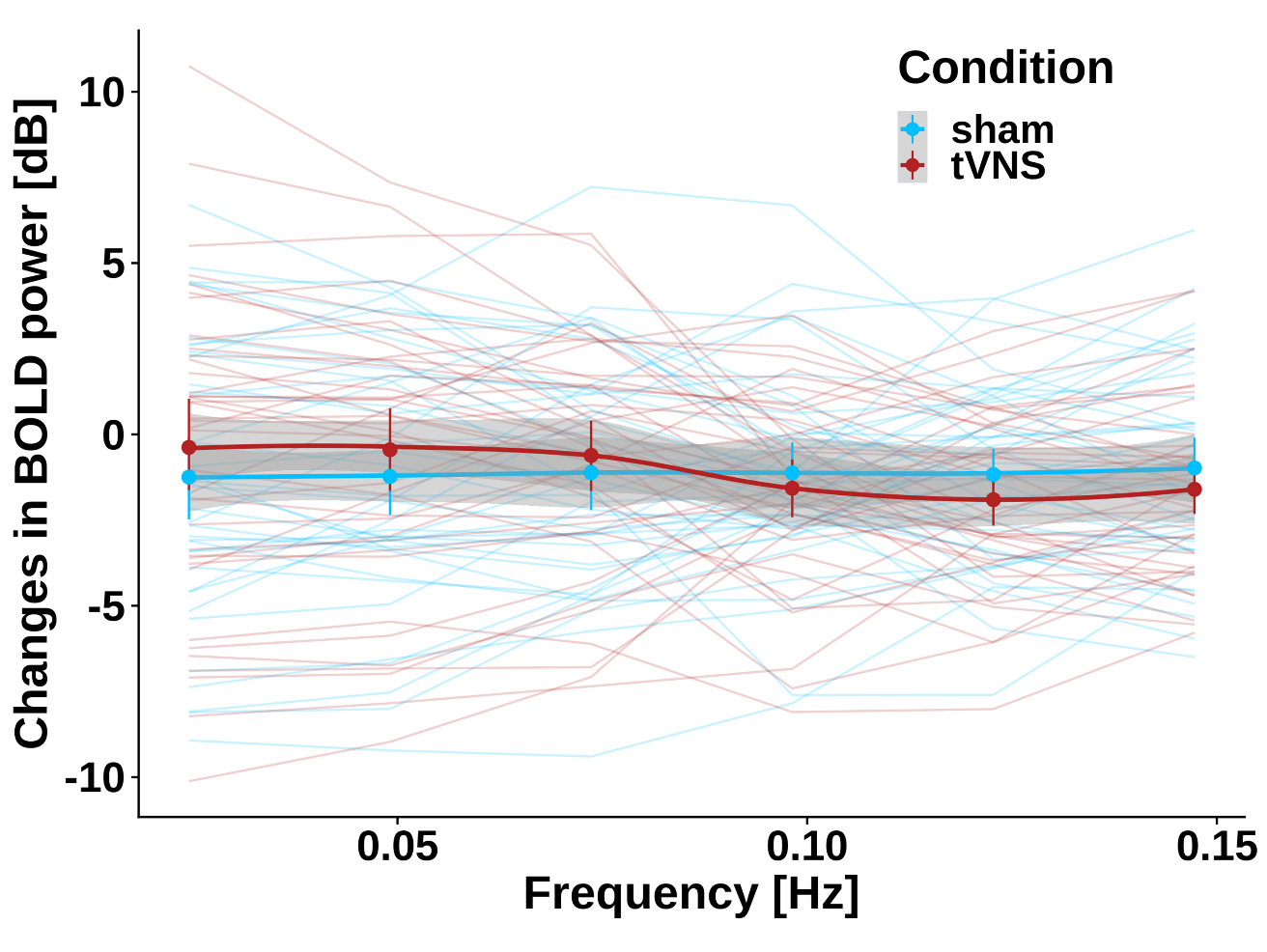

**Figure S2.** tVNS does not change the power of NTS BOLD time series compared to sham stimulation (no significant Time × Stimulation interaction). The x-axis depicts frequency bins for which we estimated power spectral density. The y-axis depicts changes in BOLD power (post - pre) for tVNS (red) and sham stimulation (blue).

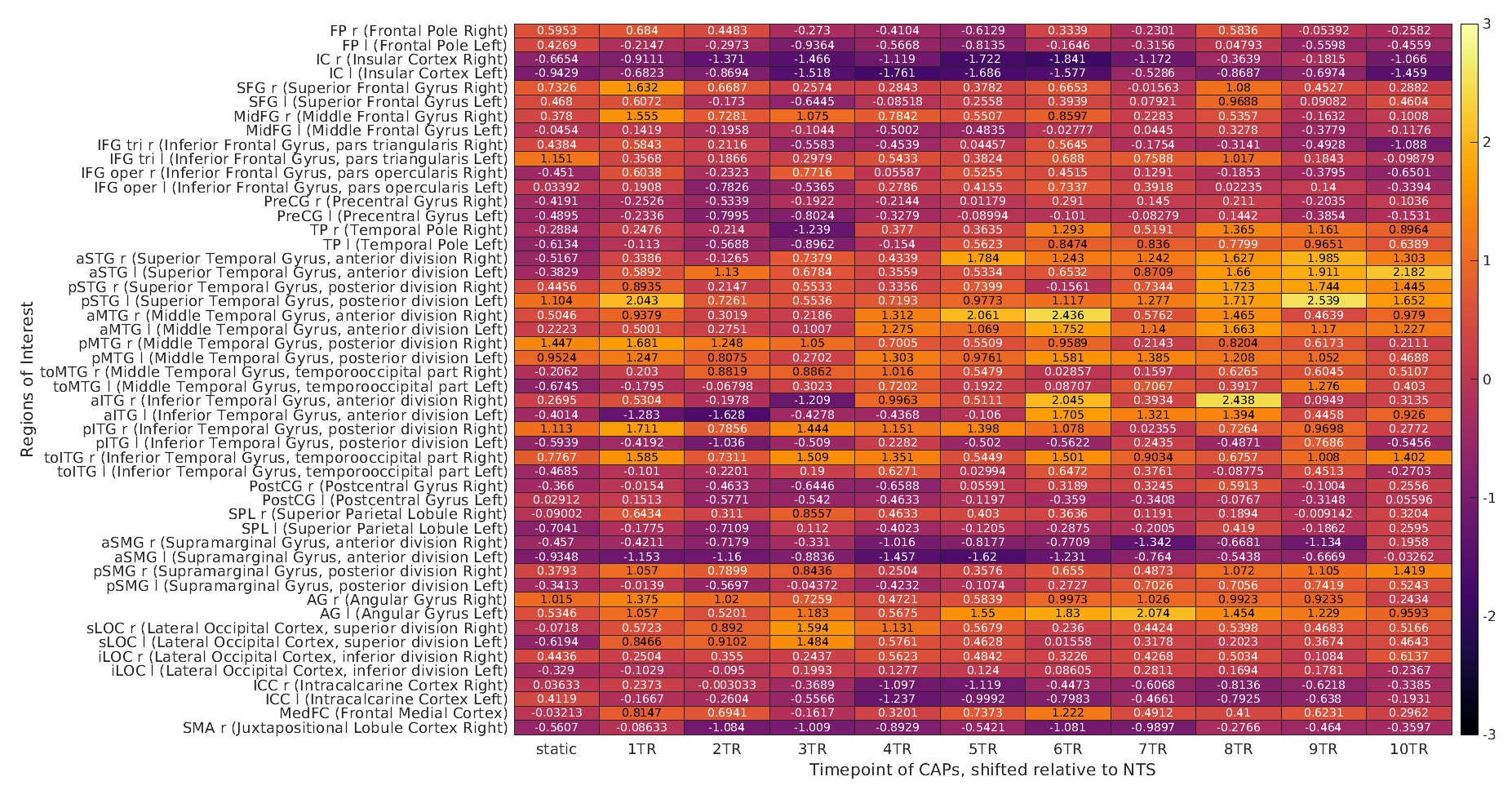

Regions of interest (Harvard-Oxford extended atlas)

Time Shift

**Figure S3.** Heatmap showing t-values for the Time (pre, post) × Stimulation (sham, tVNS) interaction predicting estimated static and dynamic co-activation patterns (CAPs) calculated between the bilateral NTS as the seed region and all regions of interest from the Harvard-Oxford extended atlas. The x-axis depicts time shifts (one TR each, ranging from static (no time shift) to a shift of 10 TR) of the NTS time series before calculating CAPs, reflecting the spatiotemporal evolution of tVNS-induced effects extending from the NTS. High positive [negative] values reflect higher activity for tVNS compared to sham [sham compared to tVNS] in the respective region on the y-axis which showed (time-lagged) co-activation with the NTS.

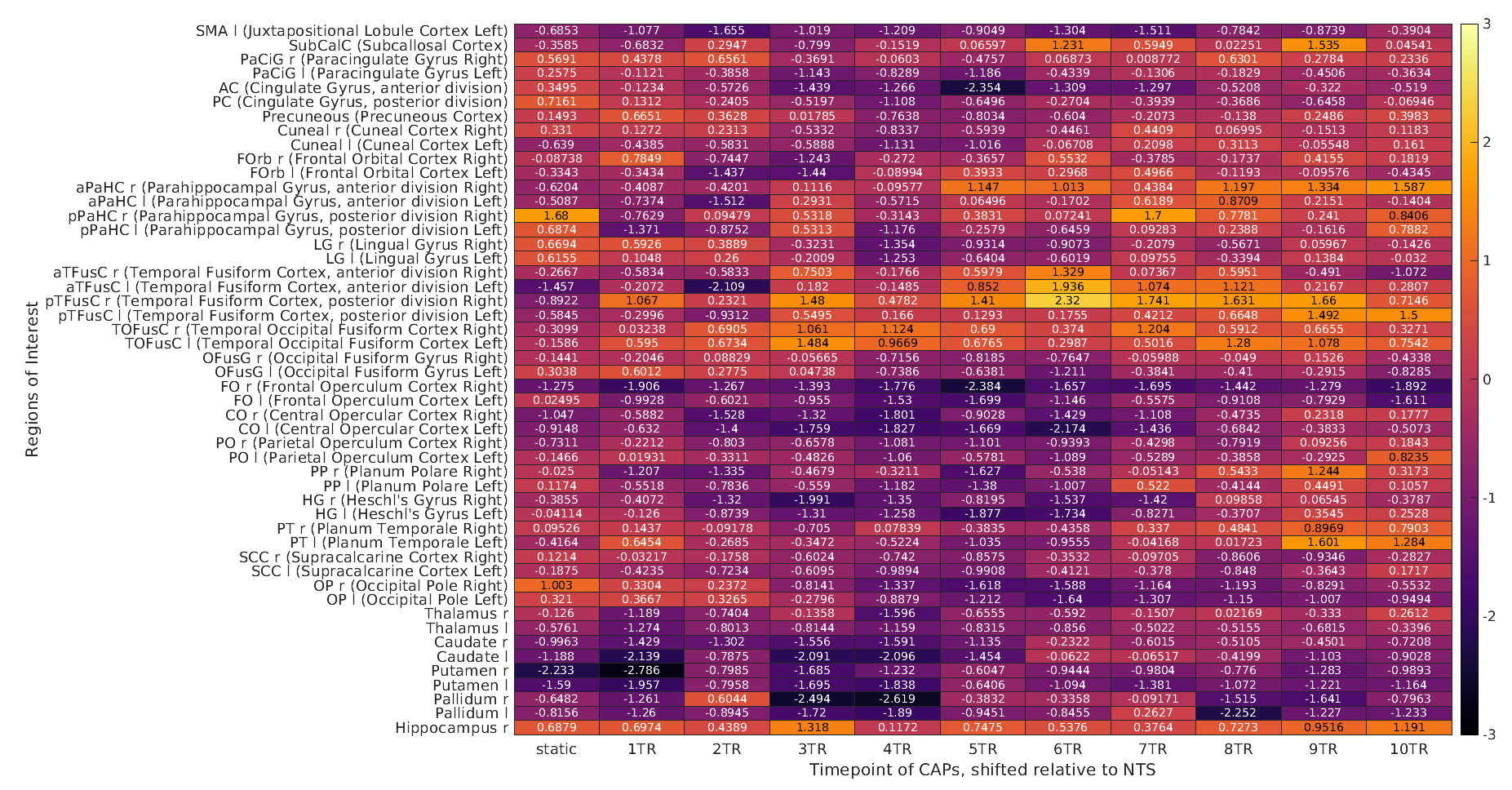

Time Shift

Regions of interest (Harvard-Oxford extended atlas)

**Figure S3 (continued).** Heatmap showing t-values for the Time (pre, post) × Stimulation (sham, tVNS) interaction predicting estimated static and dynamic co-activation patterns (CAPs) calculated between the bilateral NTS as the seed region and all regions of interest from the Harvard-Oxford extended atlas. The x-axis depicts time shifts (one TR each, ranging from static (no time shift) to a shift of 10 TR) of the NTS time series before calculating CAPs, reflecting the spatiotemporal evolution of tVNS-induced effects extending from the NTS. High positive [negative] values reflect higher activity for tVNS compared to sham [sham compared to tVNS] in the respective region on the y-axis which showed (time-lagged) co-activation with the NTS.

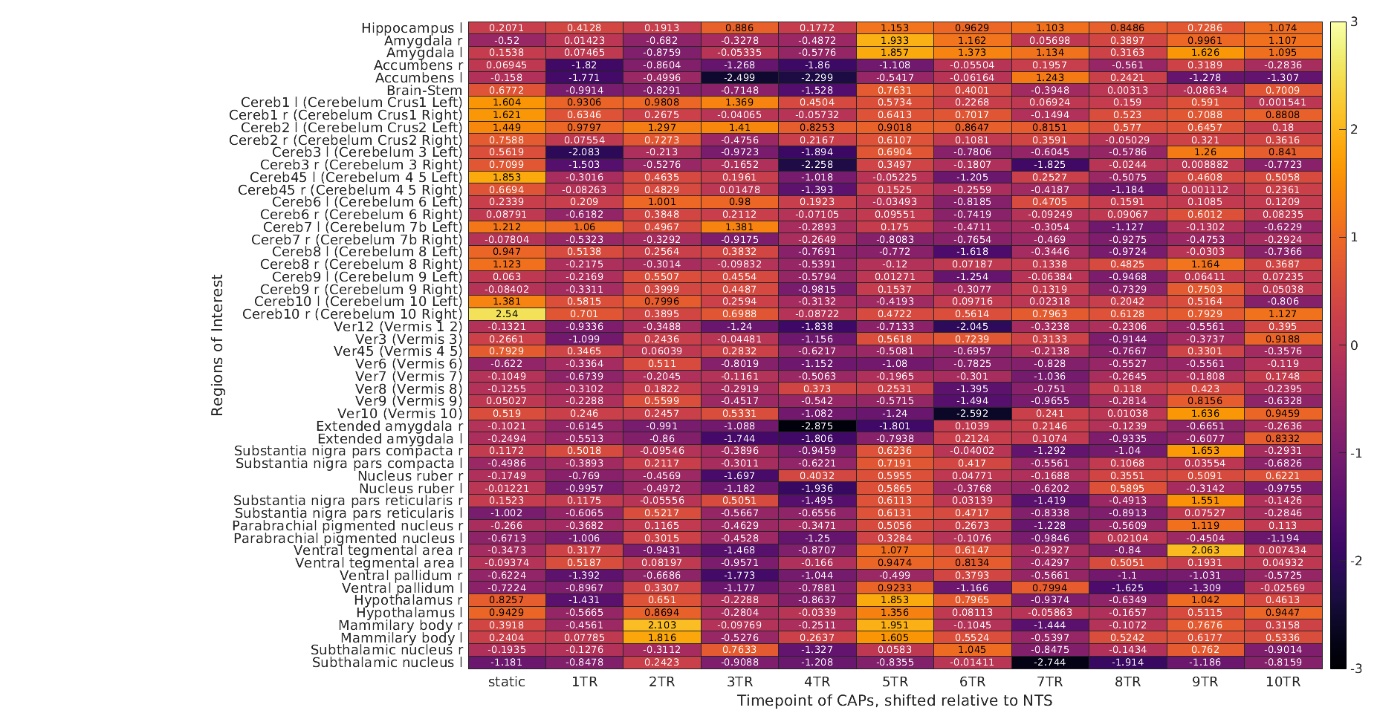

Regions of interest (Harvard-Oxford extended atlas)

Time Shift

**Figure S3 (continued).** Heatmap showing t-values for the Time (pre, post) × Stimulation (sham, tVNS) interaction predicting estimated static and dynamic co-activation patterns (CAPs) calculated between the bilateral NTS as the seed region and all regions of interest from the Harvard-Oxford extended atlas. The x-axis depicts time shifts (one TR each, ranging from static (no time shift) to a shift of 10 TR) of the NTS time series before calculating CAPs, reflecting the spatiotemporal evolution of tVNS-induced effects extending from the NTS. High positive [negative] values reflect higher activity for tVNS compared to sham [sham compared to tVNS] in the respective region on the y-axis which showed (time-lagged) co-activation with the NTS.

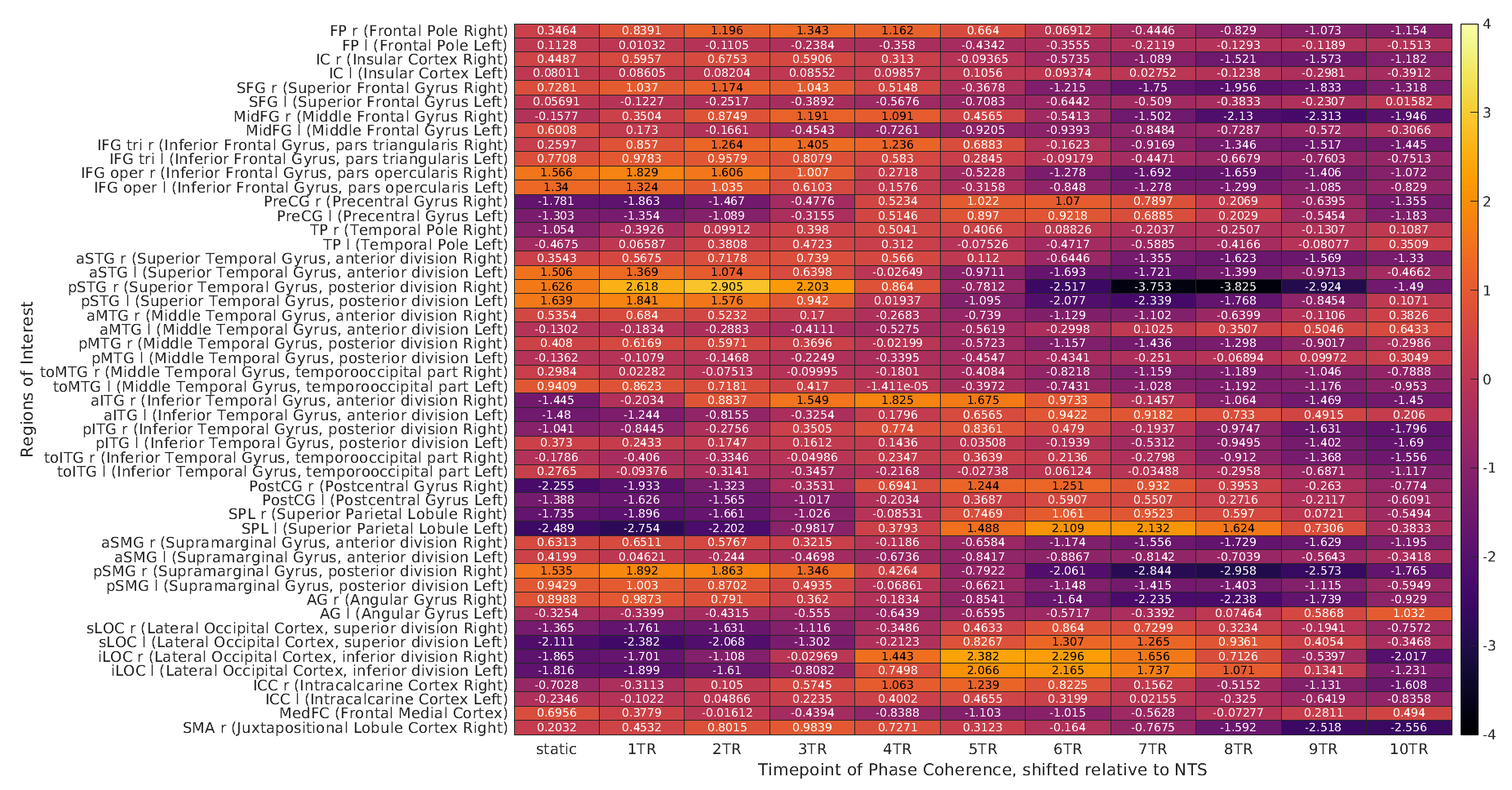

Regions of interest (Harvard-Oxford extended atlas)

Time Shift

**Figure S4.** Heatmap showing t-values for the Time (pre, post) × Stimulation (sham, tVNS) interaction predicting estimated static and dynamic phase coherence calculated between the bilateral NTS as the seed region and all regions of interest from the Harvard-Oxford extended atlas. The x-axis depicts time shifts (one TR each, ranging from static (no time shift) to a shift of 10 TR) of the NTS time series before calculating phase coherence, reflecting the spatiotemporal evolution of tVNS-induced effects extending from the NTS. High positive [negative] values reflect increased (time-lagged) phase coherence for tVNS compared to sham [sham compared to tVNS] between the respective region on the y-axis and the NTS.

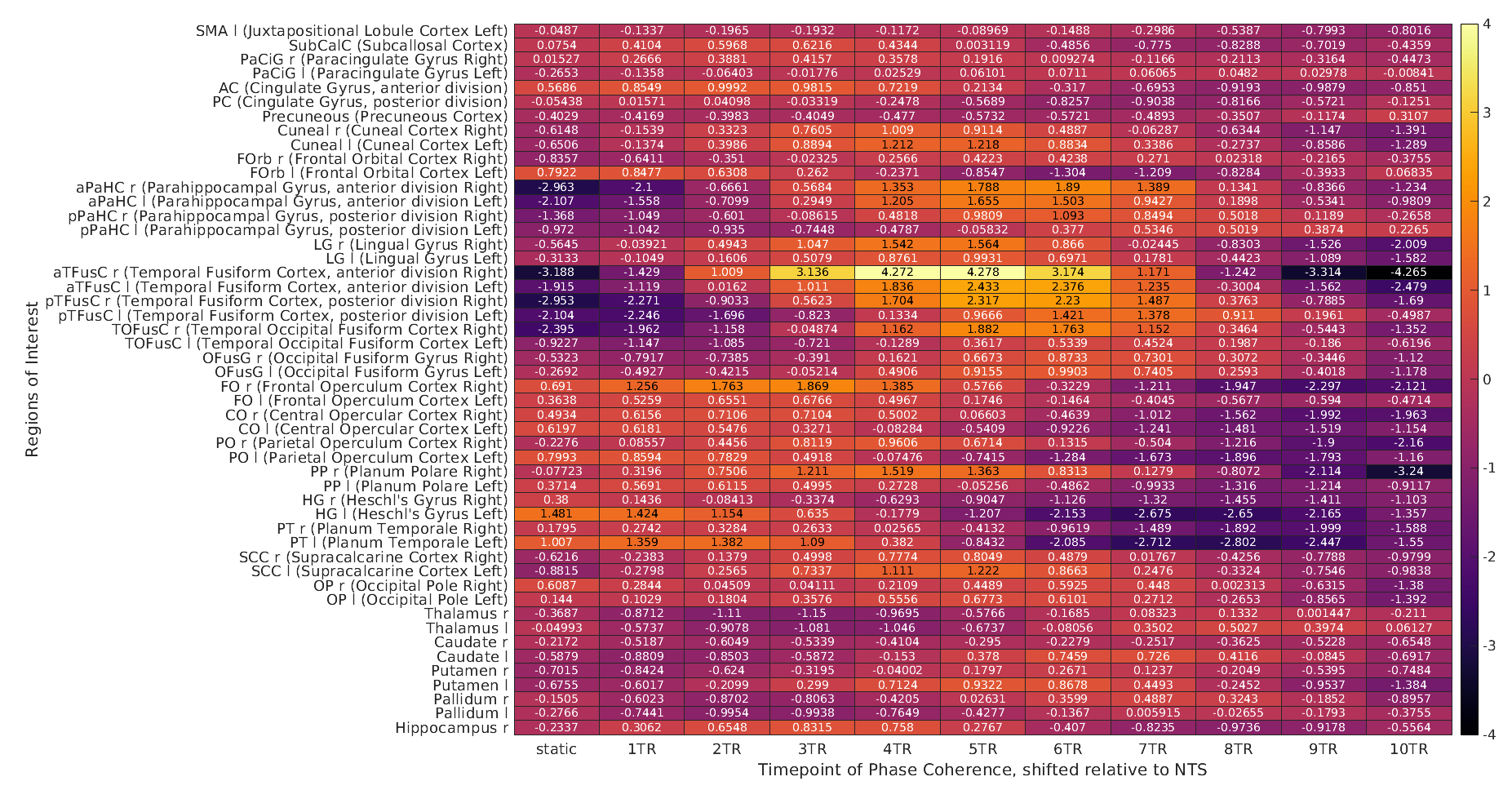

Time Shift

Regions of interest (Harvard-Oxford extended atlas)

**Figure S4 (continued).** Heatmap showing t-values for the Time (pre, post) × Stimulation (sham, tVNS) interaction predicting estimated static and dynamic phase coherence calculated between the bilateral NTS as the seed region and all regions of interest from the Harvard-Oxford extended atlas. The x-axis depicts time shifts (one TR each, ranging from static (no time shift) to a shift of 10 TR) of the NTS time series before calculating phase coherence, reflecting the spatiotemporal evolution of tVNS-induced effects extending from the NTS. High positive [negative] values reflect increased (time-lagged) phase coherence for tVNS compared to sham [sham compared to tVNS] between the respective region on the y-axis and the NTS.

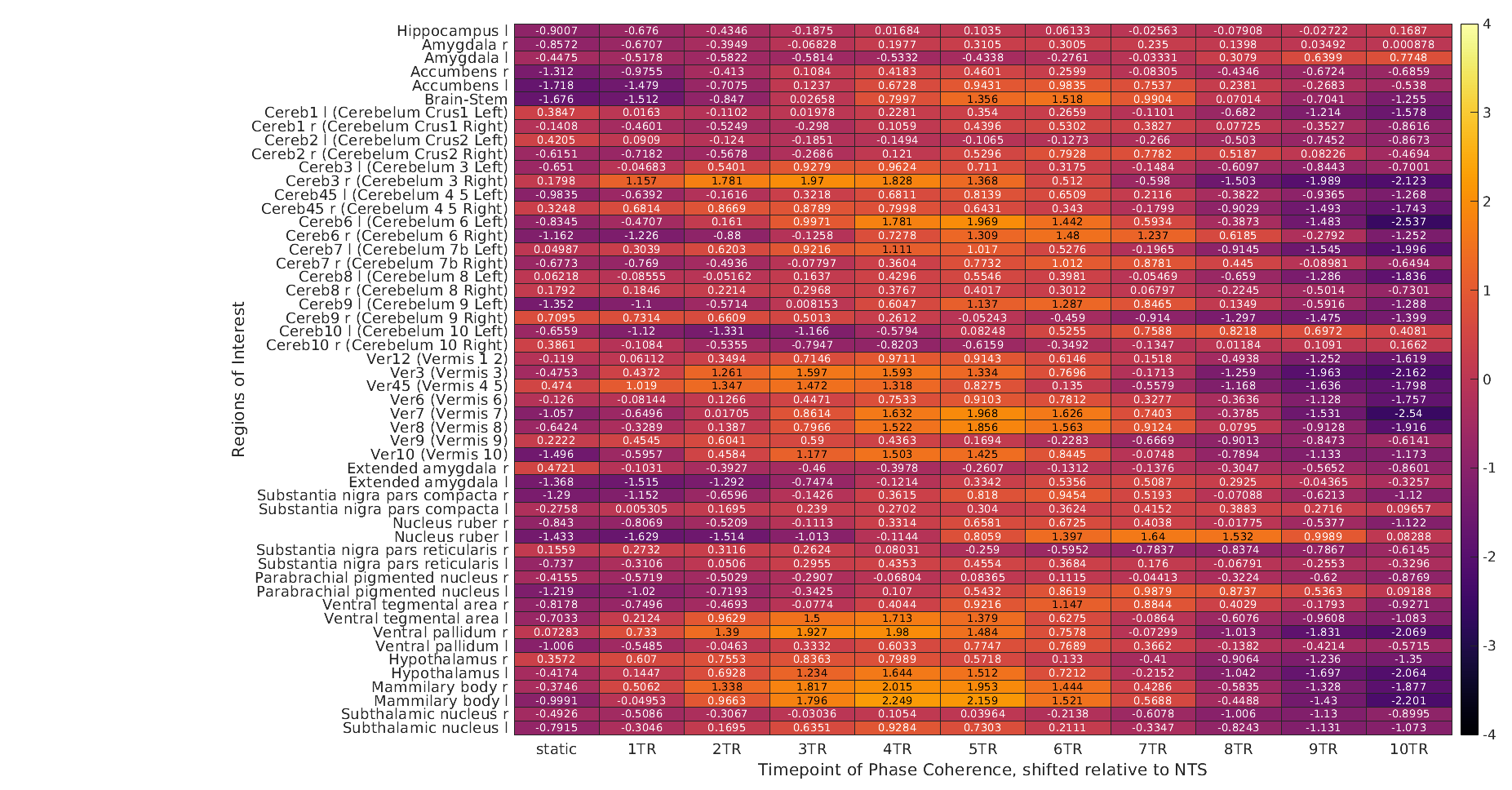

Regions of interest (Harvard-Oxford extended atlas)

Time Shift

**Figure S4 (continued).** Heatmap showing t-values for the Time (pre, post) × Stimulation (sham, tVNS) interaction predicting estimated static and dynamic phase coherence calculated between the bilateral NTS as the seed region and all regions of interest from the Harvard-Oxford extended atlas. The x-axis depicts time shifts (one TR each, ranging from static (no time shift) to a shift of 10 TR) of the NTS time series before calculating phase coherence, reflecting the spatiotemporal evolution of tVNS-induced effects extending from the NTS. High positive [negative] values reflect increased (time-lagged) phase coherence for tVNS compared to sham [sham compared to tVNS] between the respective region on the y-axis and the NTS.

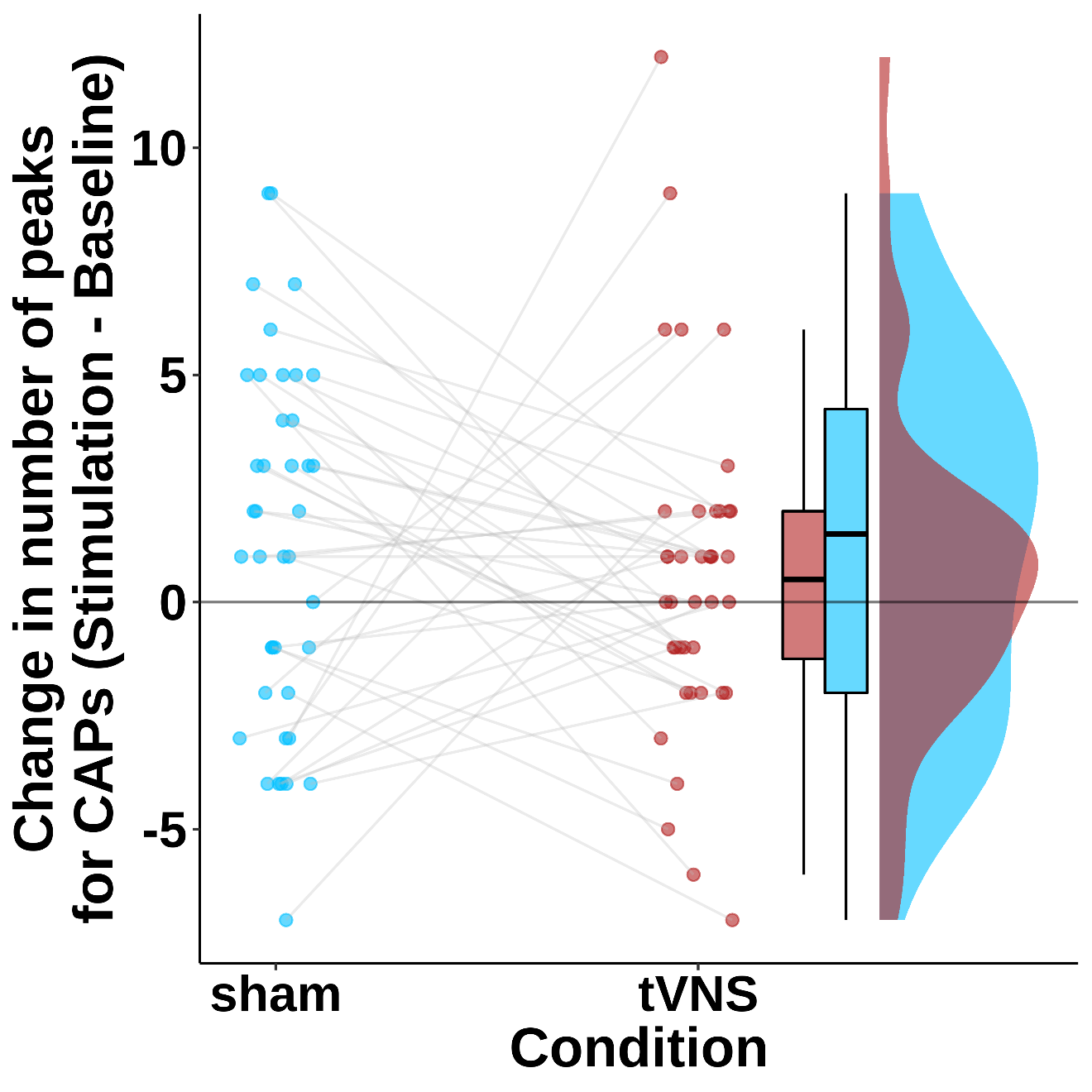

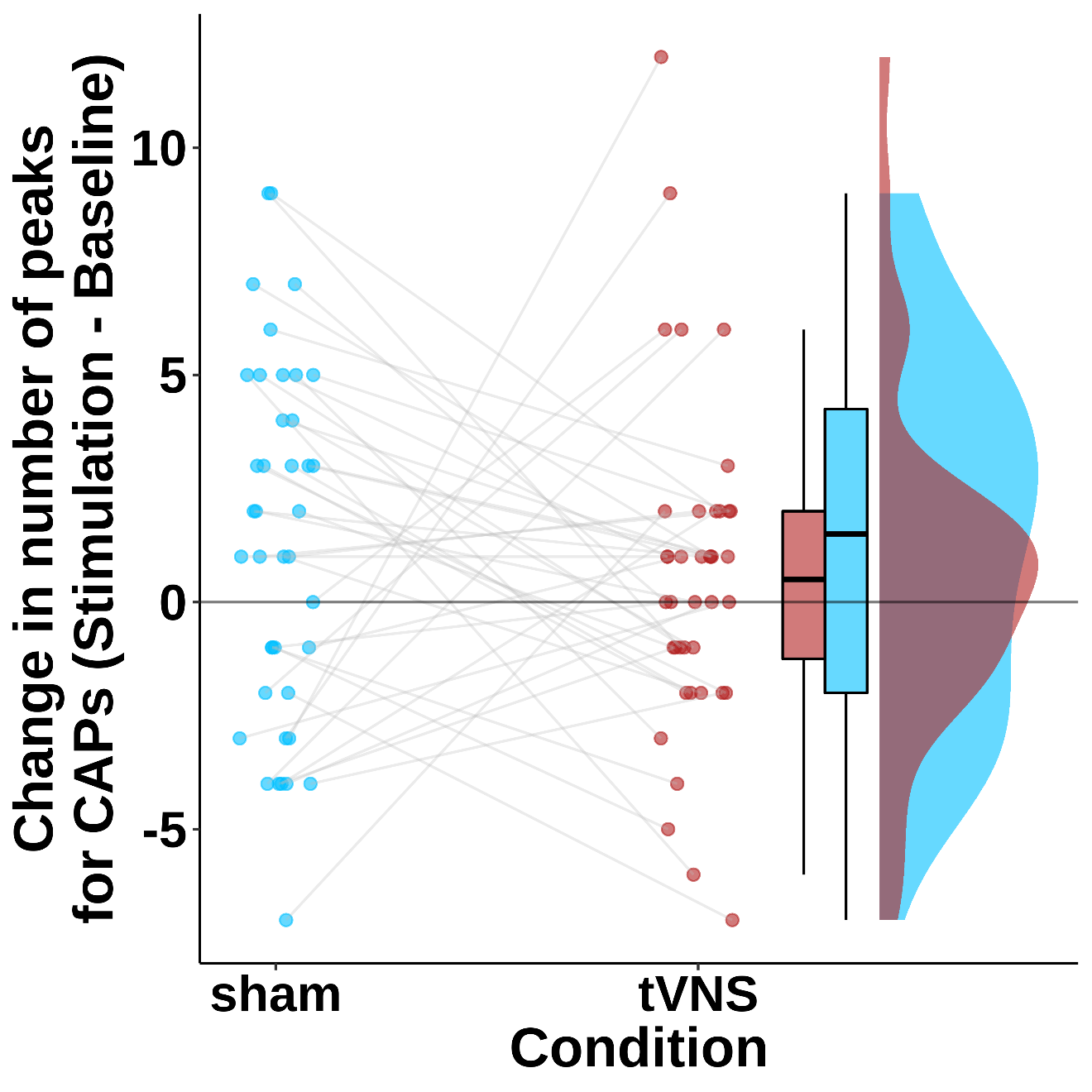

**Figure S5.** Number of peaks determined after thresholding the NTS signal for the CAP calculation does not change after tVNS compared to sham stimulation (no significant Time × Stimulation interaction). The y-axis depicts the change in number of peaks (subtracting number of peaks during baseline (pre) from the number of peaks during stimulation (post)) for tVNS (red) and sham stimulation (blue).
